# Heterogeneity in cell-cycle dynamics of synthetic mRNA-induced β-cell proliferation

**DOI:** 10.64898/2026.09.11.750925

**Authors:** Katerina Bittenglova, Klara Zacharovova, Filip Tichanek, Ivan Leontovyc, Peter Girman, Jan Kriz, Frantisek Saudek, Tomas Koblas

**Author notes:** Senior author (Tomas Koblas).

## Abstract

Adult pancreatic β-cells are mostly locked in quiescence, limiting large-scale analysis of their cell cycle. To overcome this limitation, we used synthetic *in vitro* transcribed mRNAs encoding Cyclin D1 and CDK4 to induce proliferation in up to 70% of primary rat β-cells. Flow cytometry-based analysis of cell-cycle markers revealed substantial heterogeneity in β-cell cell-cycle progression, identifying five distinct groups of proliferating β-cells based on G1 entry timing. Total cell-cycle length ranged from approximately 26 to 34 hours and was primarily determined by variability in G1 duration (13–19 hours). In the fastest-dividing β-cells, G1, S, and G2M lasted approximately 13, 6, and 7 hours, respectively, whereas later-entering β-cells exhibited progressively longer G1 phase. Additionally, we identified a small β-cell subpopulation that failed to complete division and may have exited the cell cycle. These findings reveal cell-cycle entry and G1 progression as a major source of heterogeneity in β-cell proliferation and provide quantitative benchmark for developing strategies to enhance controlled β-cell regeneration in diabetes.

## INTRODUCTION

Adult β-cells are long-lived cells with a very low proliferation rate.(Cnop *et al*, 2010) In humans, β-cells proliferation peaks during the neonatal period and declines gradually after early childhood to approximately 0.5% or less (Gregg *et al*, 2012). Physiological conditions such as pregnancy or non-diabetic obesity can induce a modest increase in β-cell mass as an adaptive response to elevated insulin demand (Butler *et al*, 2010; Saisho *et al*, 2013; Van Assche *et al*, 1978). While proliferation is a well-documented adaptive mechanism in rodents (Bock *et al*, 2003; Hull *et al*, 2005; Parsons *et al*, 1995; Rieck *et al*, 2009), expansion of β-cell mass in humans appears to rely mainly on hypertrophy and potentially neogenesis of existing β-cells, rather than on β-cell proliferation (Basile *et al*, 2019; Granger & Kushner, 2009).

Given the limited physiological proliferative capacity of β-cells, exogenous stimulation is required to activate their cell cycle and enable analysis of cell-cycle kinetics. Understanding β-cell cell-cycle dynamics may provide insight into why proliferation is normally restricted and how quiescence might be reversed. β-cell proliferation can be induced by multiple approaches, including ectopic overexpression of cell-cycle regulators (Cozar-Castellano *et al*, 2004; Fiaschi-Taesch *et al*, 2009; Guthalu Kondegowda *et al*, 2010; Tiwari *et al*, 2015), suppression of cell-cycle inhibitors (Avrahami *et al*, 2014; Robitaille *et al*, 2016; Tiwari *et al*, 2016), and, more recently, treatment with small molecules (Aamodt *et al*, 2016; Abdolazimi *et al*, 2018; Ackeifi *et al*, 2020; Shen *et al*, 2015; Wang *et al*, 2019).

Considering that the majority of adult human and rodent β-cells remain in a quiescent state (Fiaschi-Taesch *et al*, 2013a), investigating cell-cycle progression in primary β-cell cultures is challenging. Consequently, only a few studies have analysed the β-cell cell-cycle kinetics so far. These studies relied on immortalized β-cell lines (Montemurro *et al*, 2017; Andersson *et al*, 2015), partially proliferating fetal and postnatal β-cells (Swenne, 1982; Bunnag, 1966), or on induced proliferation of adult β-cells (Hija *et al*, 2014; Saisho *et al*, 2009; Ouziel-Yahalom *et al*, 2006; Russ *et al*, 2008). An early autoradiographic study of rat fetal islets *in vitro* and postnatal mouse islets *in vivo*—both predominantly composed of β-cells—estimated cell-cycle durations of approximately 14.9 hours and 10.6 hours, respectively (Swenne, 1982; Bunnag, 1966). Later studies, employing modern techniques, reported significantly longer cell-cycle durations. Juvenile rat β-cells *in vitro* required approximately 40 hours to complete the cell cycle, while adult mouse β-cells *in vivo* required 27 hours from mitogenic pulse to cytokinesis (Hija *et al*, 2014; Saisho *et al*, 2009). In the mouse study, the durations of the G1, S, and G2/M phases were estimated to be 5, 8, and 6 hours, respectively (Hija *et al*, 2014). Only one group investigated cell-cycle kinetics in human islet cells, estimating a doubling time of seven days (Ouziel-Yahalom *et al*, 2006; Russ *et al*, 2008).

A key limitation of these studies is that only a small fraction of β-cells actively proliferated. Moreover, these approaches often lacked sufficient resolution to comprehensively define cell-cycle phases in unsynchronised β-cell populations and did not allow direct confirmation of cell division.

Our laboratory has recently developed an approach that robustly activates the cell cycle in the majority of primary islet cells (Koblas *et al*, 2026). This strategy relies on ectopic overexpression of key cell-cycle regulators encoded by *in vitro* transcribed (IVT) mRNAs. The IVT mRNA-based approach has previously been used for cell fate manipulation, gene expression regulation, and gene editing with high efficiency (Koblas *et al*, 2019; Leontovyc *et al*, 2017; Warren *et al*, 2010; Gillmore *et al*, 2021). In our particular study, we used IVT mRNAs encoding cyclin-dependent kinase 4 (CDK4) and cyclin D1. Application of these IVT mRNAs resulted in a 70–80% increase in the number of β-cells after a single dose. Building on this system, we investigated cell-cycle kinetics and phase durations in primary β-cells within the islet cell cultures. Cell-cycle progression was assessed by flow cytometry (FC) using irreversible incorporation of 5-ethynyl-2′-deoxyuridine (EdU) into newly synthesized DNA combined with measurement of DNA content, allowing discrimination of nearly all cell-cycle phases (Buck *et al*, 2008). This method represents a modification of the original cell-cycle assay based on 5-bromo-2′-deoxyuridine (BrdU) incorporation (Terry & White, 2006). Unlike BrdU, EdU detection does not require harsh DNA denaturation and is now widely used for analysis of cell-cycle progression, particularly in oncology studies (Jolly *et al*, 2022; Frölich *et al*, 2020b).

Here, we provide a high-resolution characterisation of asynchronous and partially synchronised populations of proliferating β-cells across different cell-cycle phases. We show that proliferating β-cells display marked heterogeneity in the timing of G1 entry, cell-cycle kinetics, and subsequent cell-cycle progression. Based on the G1 entry timing, we identified five distinct groups of proliferating β-cells with different kinetics, revealing that later G1 entry is associated with progressively longer cell-cycle duration. These observations suggest that differences in G1 duration largely determine variability in β-cell cell-cycle length. Furthermore, we established a combination of cell-cycle markers enabling detailed analysis of progression through individual cell-cycle phases.

## RESULTS

### Half of β-cells complete division within 40 hours after IVT mRNA-induced cell-cycle activation

With the aim to conduct detailed study of β-cell proliferation, we employed optimised flow cytometry to analyse cell-cycle progression and detect effective cell division. This method combines irreversible EdU incorporation into replicating DNA with measurement of total DNA content to resolve distinct cell-cycle states during proliferation. Based on the EdU intensity and DNA content, proliferating cells can be divided into the following subpopulations. The G0/G1 EdU^-^cell population with 2N DNA content corresponded to quiescent G0 or G1 phase. S-phase cells (S EdU^+^) incorporated EdU and exhibited intermediate DNA content (2N–4N), whereas G2/M EdU^+^ cells displayed maximal EdU intensity and 4N DNA content. After cytokinesis/cell division, daughter cells regained 2N DNA content while retaining approximately half of the incorporated EdU signal from the mother cell. Cells entering a second round of the cell cycle (2nd S) exhibited increasing DNA and EdU fluorescence while retaining EdU label from the previous round. A minor G2/M EdU^-^population (unlabelled 4N cells) was also detectable but remained negligible under non-proliferative conditions.

To enable precise temporal analysis of β-cell cell-cycle progression, proliferating islet cells were sampled at 2-hour intervals during the first 34 hours post-transfection (hpt) of IVT mRNAs encoding Cyclin D1 and CDK4, and subsequently every 10 hours up to 70 hpt, as outlined in the experimental timeline in Fig 1A.

**Fig 1:**
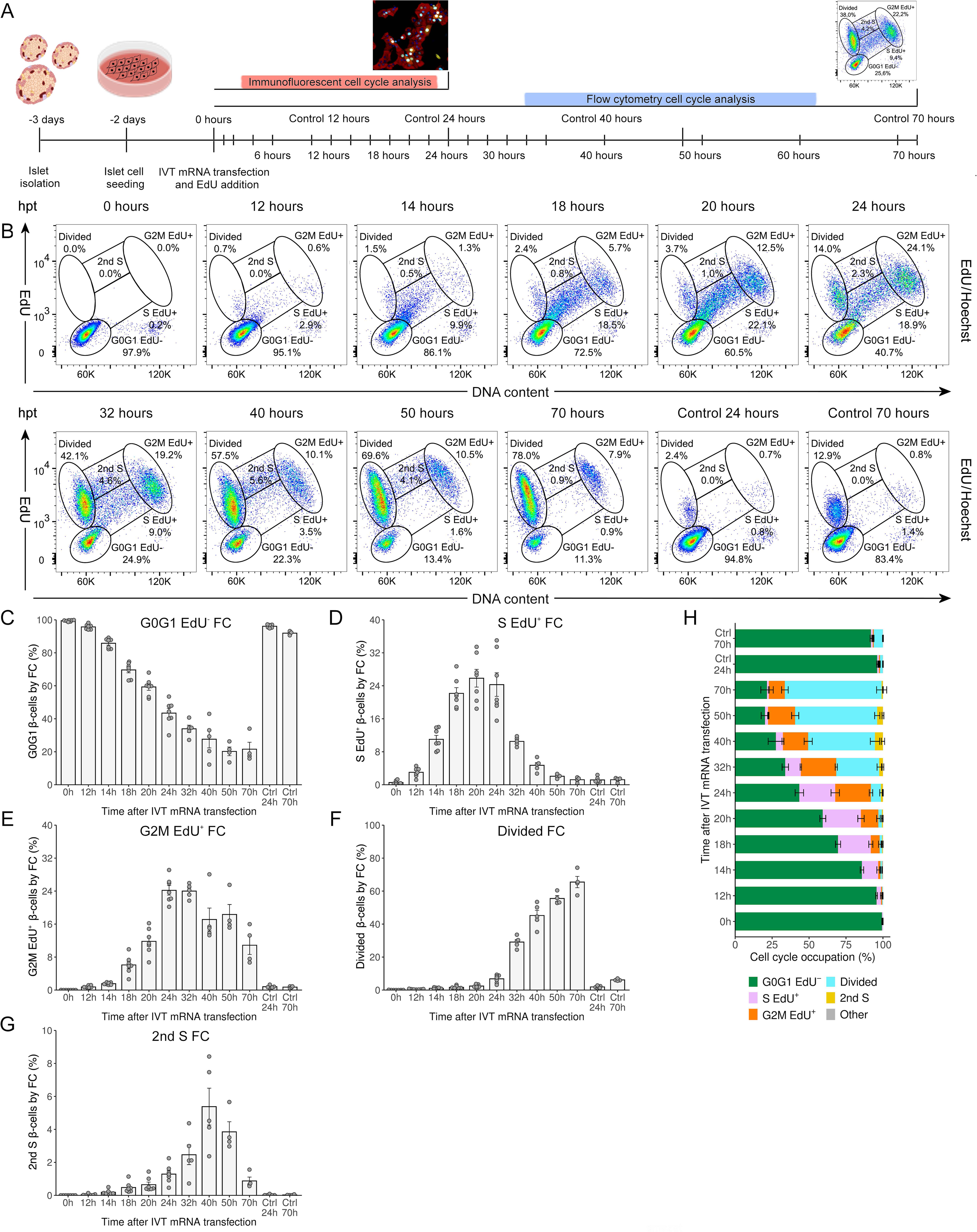
Half of β-cells complete division within 40 hours after IVT mRNA-induced cell-cycle activation (A) Schematic timeline illustrating the experimental design (see also Methods). (B) Representative flow cytometry (FC) plots of proliferating β-cells showing EdU incorporation and DNA content at selected time points compared to untreated controls (see also Fig EV1). (C–G) Proportions of β-cells in G0/G1 EdU^-^(C), S EdU^+^ (D), G2/M EdU^+^ (E), divided (F), and 2nd S (G) quantified from EdU/DNA flow cytometry plots and normalized to the original number of β-cells at the same time points as in (B) (n=4–7 biological replicates). (H) Distribution of β-cells across cell-cycle phases derived from the quantifications shown in (C–G) and remaining cells grouped as “Other”. Bars represent mean ± SEM (C–H). Omnibus tests for the overall effect of experimental group (time point or control) were *P* < 0.001 for all tested models (C–G), based on linear mixed-effects models with experiment included as a random intercept, experimental group included as a categorical predictor, and group-specific residual variances allowed. Complete pairwise comparison results are provided in the online statistical report described in the Data and code availability statement. Hpt, hours post-transfection

Using insulin staining to distinguish β-cells from other islet cell types, we found that the proportion of divided β-cells increased progressively and reached 78.07 ± 2.41% at 70 hpt, compared with 11.39 ± 0.53% in the untreated control (Fig 1B and EV1). At the same time point, the overall fraction of EdU^+^ β-cells approached 87.09 ± 2.72%, indicating robust induction of cell-cycle entry. Nonetheless, most β-cells divided substantially earlier than 70 hpt. The first divided β-cells were detected already at 26 hpt (20.02 ± 2.17%), followed by an approximately twofold increase by 30 hpt (39.01 ± 2.15%). By 40 hpt, more than half of the β-cells had completed at least one division. However, once cell division commenced, population frequencies no longer reflected the fraction of the original β-cell population because cytokinesis doubled the number of divided cells. To determine the fraction of the original β-cells that completed division, we recalculated population frequencies derived from EdU/DNA flow cytometry plots using the normalization equations described in the Methods. Following this correction, the values changed slightly: the absolute proliferation efficiency of the original β-cell population reached 65.48 ± 3.42% at 70 hpt while the remaining β-cells failed to complete division during the monitored interval (Fig 1F and 1I). In contrast, 21.56 ± 4.20% of β-cells exhibited quiescence throughout the experiment (Fig 1C and 1I), while 10.87 ± 2.24% remained arrested in G2/M without completing division (Fig 1E and 1I). The persistent presence of this G2/M EdU^+^ population at 60 to 70 hpt may reflect cell-cycle exit or senescence, although we cannot rule out delayed division associated with prolonged G2/M. Consistent with this, the proportion of Cyclin A2-negative β-cells within the G2/M population increased between 40 and 70 hpt (Fig EV2), together with a decrease in phosphorylated retinoblastoma protein (pRB) positivity (Fig EV3). During physiological mitosis, Cyclin A2 expression and RB phosphorylation are rapidly lost (Ludlow *et al*, 1993; Den Elzen & Pines, 2001). Loss of Cyclin A2 during G2/M was linked to APC/C^Cdh1^-mediated degradation during the cell-cycle exit, a process associated with senescence detectable after 2-3 days (Wiebusch & Hagemeier, 2010; Johmura *et al*, 2014; Müllers *et al*, 2014). The elevated G2/M fraction was unlikely to be explained by continued entry of β-cells into G2/M, as only a few β-cells remained in the S-phase beyond 40 hpt (Fig 1B, 1D, 1G, and S1). Additionally, we detected a small population of EdU^-^4N β-cells at 0 hpt that persisted throughout the whole experiment. However, its size decreased over time, suggesting either apoptotic loss or delayed mitotic completion.

Interestingly, we observed the emergence of a new population of divided β-cells undergoing a second round of DNA replication from approximately 30 hpt. These cells showed higher EdU fluorescence intensity than β-cells in the first cell cycle (Fig 1B, 1G, 1I, and S1). Entry into the second cell cycle was further supported by the reappearance of pRB positivity (Fig EV1 and EV3). Consistently, the median of EdU intensity within the divided β-cell population gradually increased over time, consistent with repeated DNA synthesis and progression through a second cell cycle. The induction of a second cell cycle coincided with continuous exposure to IVT mRNAs during the first 48 hpt, after which the medium was replaced.

However, the emergence of a second cell cycle introduced a major limitation for kinetic analyses, as the G2/M and divided populations became a mixture of β-cells belonging to either the first or the second cell cycle. Consequently, it was no longer possible to unambiguously distinguish between these two rounds of division within these subpopulations. Therefore, detailed analyses of β-cell cell-cycle kinetics were restricted to approximately the first 34 hpt.

Taken together, IVT mRNA-induced cell-cycle entry did not synchronise β-cells across cell-cycle phases, resulting in a heterogeneous proliferative progression. To further characterise this heterogeneity of β-cell-cycle progression, we next examined the timing of G1 entry by monitoring RB phosphorylation.

### β-cell G1 duration varies with the timing of G1 entry

We monitored G1 entry by assessing phosphorylation of the retinoblastoma protein (RB), a central gatekeeper of G1 entry (Weinberg, 1995). Hypophosphorylated RB suppresses E2F transcription factors’ activity and cell-cycle progression, whereas RB phosphorylation by CDK4/6–cyclin D complexes results in E2F activation and promotes G1 progression (Hiebert *et al*, 1992; Dick & Rubin, 2013). We used RB phosphorylation at Ser807/811 (pRB) as a marker of G1 entry based on its previous use in rat β-cells (Tiwari *et al*, 2015).

As shown in Fig 2B and 2D–2E, the proportion of pRB^+^ β-cells increased rapidly after stimulation, reaching 15.79 ± 2.04% at 1 hpt, 26.17 ± 3.01% at 2 hpt, and 40.89 ± 2.19% at 4 hpt. The rate of cell-cycle entry slowed thereafter, reaching approximately half of all β-cells in G1 by 6 hpt and 64.54 ± 3.22% by 12 hpt. By 24 hpt, the proportion of β-cells in G1 increased modestly to 69.32 ± 1.45% (Fig EV3). Although pRB was not quantified in detail at later time points, the fraction of EdU^-^β-cells at 70 hpt (21.56 ± 4.20%) suggests that only a small subset of β-cells (approximately 9%) entered the cell cycle after 24 hpt.

**Fig 2:**
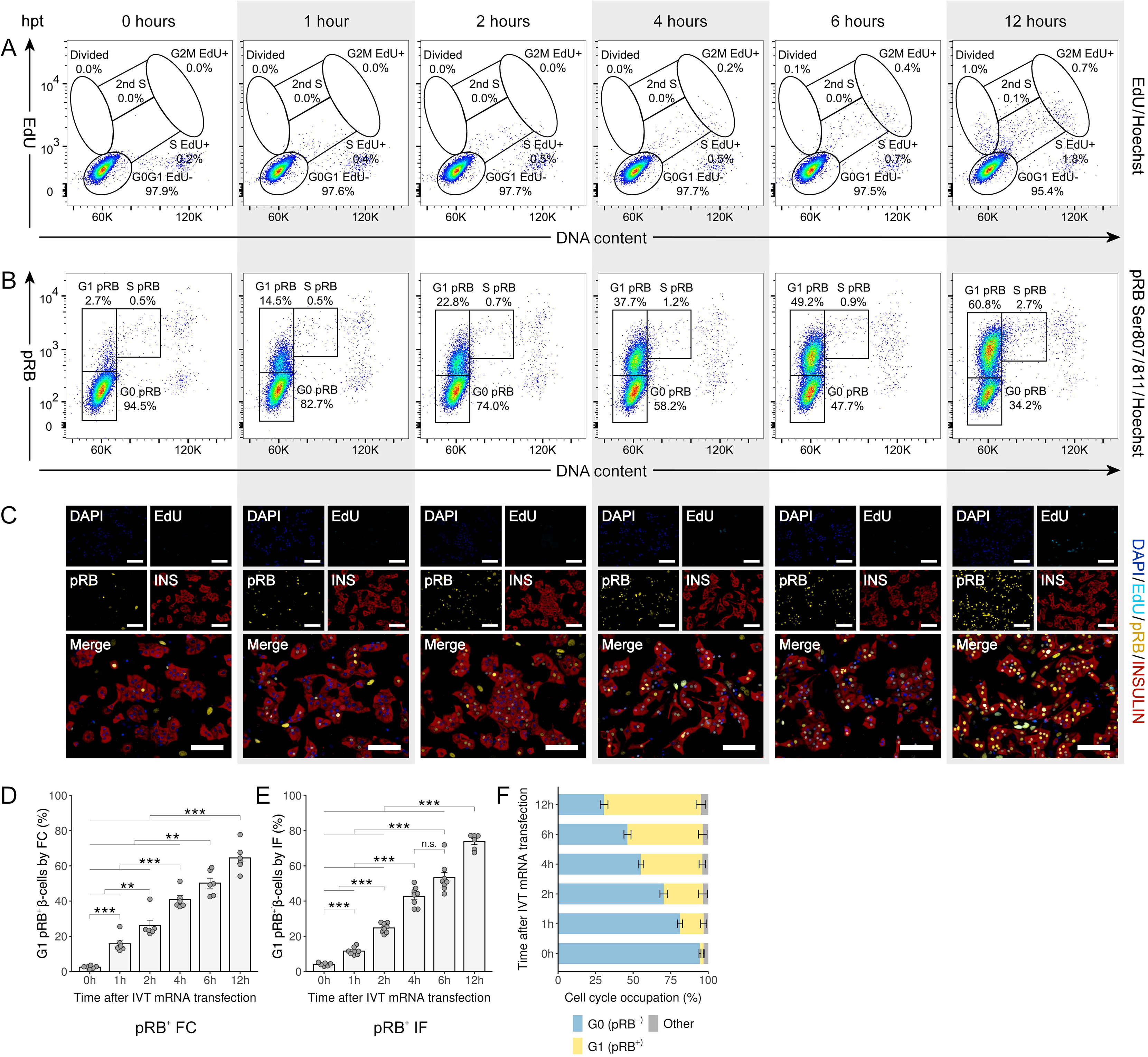
β-cell G1 duration varies with the timing of G1 entry Representative flow cytometry and immunofluorescence images at selected time points (A–C). The same time points are highlighted by white and gray shading across panels. (A–B) Representative flow cytometry plots of mRNA-treated β-cells showing EdU incorporation (A) or RB phosphorylation (pRB; B) and DNA content, compared with untreated control (see also Fig EV1 and EV3). (C) Representative immunofluorescence images of proliferating β-cells showing EdU incorporation (light blue) and immunostaining for pRB (yellow), insulin (red), and DAPI (dark blue). Scale bars, 100 µm. (D) Proportion of pRB^+^ β-cells in G1 quantified from pRB/DNA flow cytometry (FC) plots (n=6 biological replicates). (E) Proportion of pRB^+^ β-cells in G1 quantified by immunofluorescence (IF) and high-content imaging analysis (n=6–8 biological replicates). (F) Distribution of β-cells across cell-cycle phases derived from pRB/DNA flow cytometry quantification, with remaining cells grouped as “Other”. Bars represent mean ± SEM (D–F). Statistical significance was assessed using a linear mixed-effects model, including experiment as a random intercept, time as a categorical predictor (D, E), and allowing for group-specific residual variance (E). \*\*\**P* < 0.001, \*\**P* < 0.01, and \**P* < 0.05, n.s., not significant (Bonferroni-adjusted pairwise differences; omnibus test *P* < 0.001 for all tested models). Hpt, hours post-transfection

Notably, the maximal degree of RB phosphorylation was comparable between stimulated and non-stimulated β-cells, indicating that IVT mRNA-induced cell-cycle entry did not cause supraphysiological RB phosphorylation but rather recapitulated a physiologically regulated pattern of G1 activation (Fig EV3).

Based on the timing of first pRB acquisition, we stratified proliferating β-cells into five groups. The fastest cycling β-cells (group 1; 13.31 ± 2.28%) entered G1 before 1 hpt, followed by group 2 (10.38 ± 1.20%) entering G1 between 1 and 2 hpt, group 3 (14.72 ± 1.11%) between 2 and 4 hpt, group 4 (9.30 ± 2.37%) between 4 and 6 hpt, and the slowest proliferating group 5 (10.06 ± 2.31%) between 6 and 10 hpt. We excluded β-cells entering G1 later from detailed kinetic analyses as discussed above. We maintained this grouping throughout subsequent analyses. Population sizes are reported either directly from flow cytometry plots or after normalization to the original β-cell number for EdU-based analyses (Tables S1 and S2). For example, 23.69 ± 3.24% pRB^+^ β-cells at 2 hpt comprise 13.31 ± 2.28% of β-cells that had already entered G1 by 1 hpt (group 1) and 10.38 ± 1.20% that entered G1 between 1 and 2 hpt (group 2). We interpolated transitions between cell-cycle phases occurring between sampling intervals as described in the Methods.

We defined G1 duration as the interval between the onset of RB phosphorylation in a given β-cell group and the subsequent onset of EdU incorporation, marking entry into S-phase. Following the peak of pRB positivity (Fig 2B and 2D), β-cells progressively transitioned into early S-phase characterised by EdU incorporation and increasing DNA content (Fig 2A, 3A, 3D, and 3F). The earliest β-cell group 1 completed G1 phase at 14 hpt, corresponding to a G1 duration of approximately 13 hours. Group 2 transitioned into S-phase by 18 hpt, corresponding to a G1 duration of approximately 16 hours. Similarly, group 3 became EdU^+^ around 20 hpt, indicating the same G1 length. For group 4 and group 5, entry into S-phase occurred before 23 hpt and 29 hpt, corresponding to G1 durations of approximately 17 hours and up to 19 hours, respectively (Table S1).

Collectively, these analyses revealed that β-cells entered G1 asynchronously following IVT mRNA stimulation. Importantly, the duration of G1 was shortest for the earliest-entering β-cells and progressively increased in β-cells entering G1 later.

To validate these findings using an alternative approach, we analysed RB phosphorylation by immunofluorescence microscopy (IF) (Fig 2C and 2E). Weak pRB positivity was detectable in a subset of β-cells as early as 1 hpt (11.61 ± 0.72%) and increased by 2 hpt (24.78 ± 0.96%), although signal intensity at these early stages was low. Over time, both the fraction of pRB^+^ β-cells and signal intensity increased in a pattern comparable to that observed by flow cytometry. Overall, although IF showed similar trends, early weak RB phosphorylation was less unambiguous due to background signal, highlighting the higher sensitivity of flow cytometry for resolving early G1 entry.

### β-cell S-phase duration is independent of the timing of cell-cycle entry

To monitor S-phase progression, we primarily used EdU incorporation combined with DNA content analysis, which enabled clear delineation of S-phase boundaries by flow cytometry (Fig 3A, 3D, 4A, 4D, and S1). To further support S-phase identification, we analysed Cyclin A2 and stem loop binding protein (SLBP), whose expression changes during late G1 and early S-phase (Fig 3B–3C, 3H, and 3J) (Zwicker *et al*, 1995; Whitfield *et al*, 2000). However, because expression of these markers partially overlaps at phase boundaries, we did not use them to discriminate adjacent cell-cycle phases precisely.

**Fig 3:**
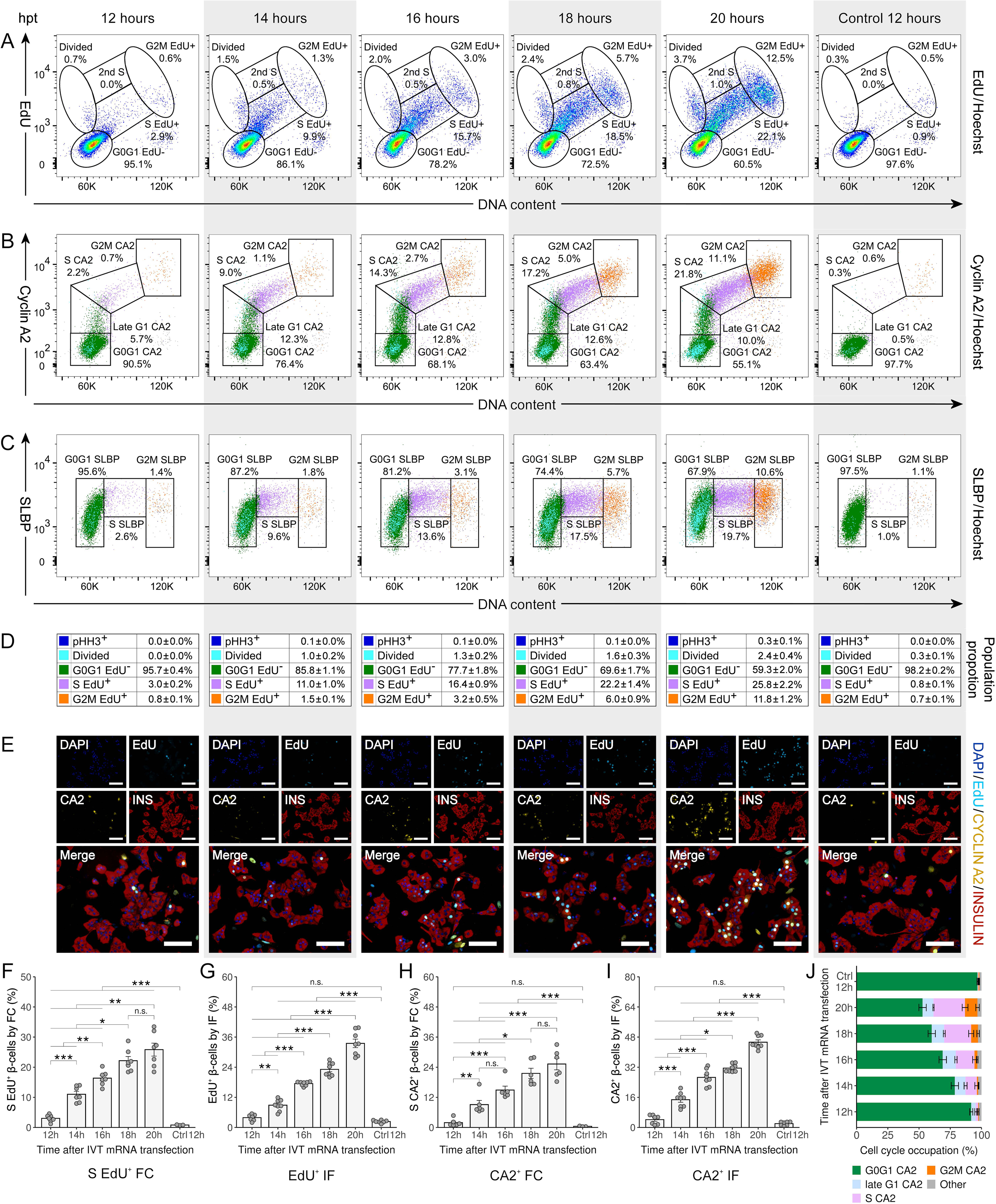
β-cell S-phase duration is independent of the timing of cell-cycle entry Representative flow cytometry plots and immunofluorescence images illustrating S-phase progression in mRNA-treated and untreated β-cells at selected time points, highlighted by matching white and gray shading across panels (A–E). Cell populations are colour-coded in panels (B–D) and in distribution graph of β-cells across cell-cycle phases (J) as follows: G0/G1 EdU⁻ (dark green), S EdU^+^ (violet), G2/M EdU^+^ (orange), divided β-cells (light blue), and pHH3^+^ (phosphorylated histone H3; dark blue). (A–C) Representative flow cytometry plots of β-cells showing EdU incorporation (A), Cyclin A2 (CA2; B), or SLBP expression (C) in combination with DNA content (see also Fig EV1 and EV2). (D) Summary table showing percentages of β-cell populations quantified from normalized EdU/DNA and pHH3/DNA flow cytometry analyses (mean ± SEM). (E) Representative immunofluorescence images of proliferating β-cells labelled with EdU (light blue) and stained for Cyclin A2 (yellow), insulin (red), and DAPI (dark blue). Scale bars, 100 µm. (F) Proportion of S-phase EdU^+^ β-cells quantified from EdU/DNA flow cytometry (FC) plots and normalized to the original β-cell population (n=3–7 biological replicates). (G) Proportion of EdU^+^ β-cells quantified by immunofluorescence (IF) and high-content imaging analysis (n=7 biological replicates). (H) Proportion of S-phase Cyclin A2^+^ β-cells quantified from Cyclin A2/DNA flow cytometry plots (n=3–7 biological replicates). (I) Proportion of Cyclin A2^+^ β-cells quantified by immunofluorescence and high-content imaging analysis (n=7 biological replicates). (J) Distribution of β-cells across cell-cycle phases derived from Cyclin A2/DNA flow cytometry quantification, with remaining cells grouped as “Other”. Bars represent mean ± SEM (F–J). Statistical significance was assessed using a linear mixed-effects model, including experiment as a random intercept, time as a categorical predictor (F–I), and allowing for group-specific residual variances (F). \*\*\**P* < 0.001, \*\**P* < 0.01, \**P* < 0.05, and n.s., not significant (Bonferroni-adjusted pairwise differences; omnibus test *P* < 0.001 for all tested models). Hpt, hours post-transfection

As described above, the first significant increase in the S-phase EdU^+^ β-cell population was detected at 14 hpt and corresponded to group 1 (13.31 ± 2.28% of the original β-cells) (Fig 3A, 3D, and 3F). The proportion of S-phase β-cells gradually increased until approximately 20 hpt, reflecting ongoing DNA replication (Fig 4A, 4D, 4F, and EV1). Cyclin A2 expression became detectable in late G1 at 12 hpt (Fig 3B, 3H) and preceded DNA replication, consistent with its role in replication control and prevention of re-licensing during S-phase.(Petersen *et al*, 1999; Bendris *et al*, 2012; Sugimoto *et al*, 2004) In parallel, SLBP expression increased during G1 and peaked at the onset of S-phase (Fig 3C), consistent with its role in histone biosynthesis during DNA replication.(Whitfield *et al*, 2000) S-phase duration was defined as the interval between the onset of EdU incorporation in a given β-cell group (Fig 3D and 3F) and its subsequent entry into the G2/M EdU^+^ gate, characterised by maximal EdU intensity and 4N DNA content (Fig 4D, 4F, and 4H). Using this approach, the most rapidly dividing β-cells (group 1, 13.31 ± 2.28%) spent approximately 6 hours in S-phase, spanning from 14 to 20 hpt. Slightly delayed β-cell group 2, representing 10.38 ± 1.20% of the population (23.69 ± 3.24% in cumulative plots), entered S-phase before 18 hpt and transitioned to G2/M by 23 hpt, indicating an S-phase duration of approximately 5 hours. Groups 3 (38.41 ± 2.41%) and 4 (47.71 ± 3.00%) exhibited similar or slightly longer S-phases lasting approximately 7 and 6 hours, respectively (Table S1). Precise determination of S-phase length in group 5 was not possible because consecutive cell cycles overlapped, as discussed above. Overall, the S-phase duration remained relatively uniform across β-cell subpopulations, ranging from approximately 5 to 7 hours and, unlike G1 duration, did not correlate with the timing of cell-cycle entry.

**Fig 4:**
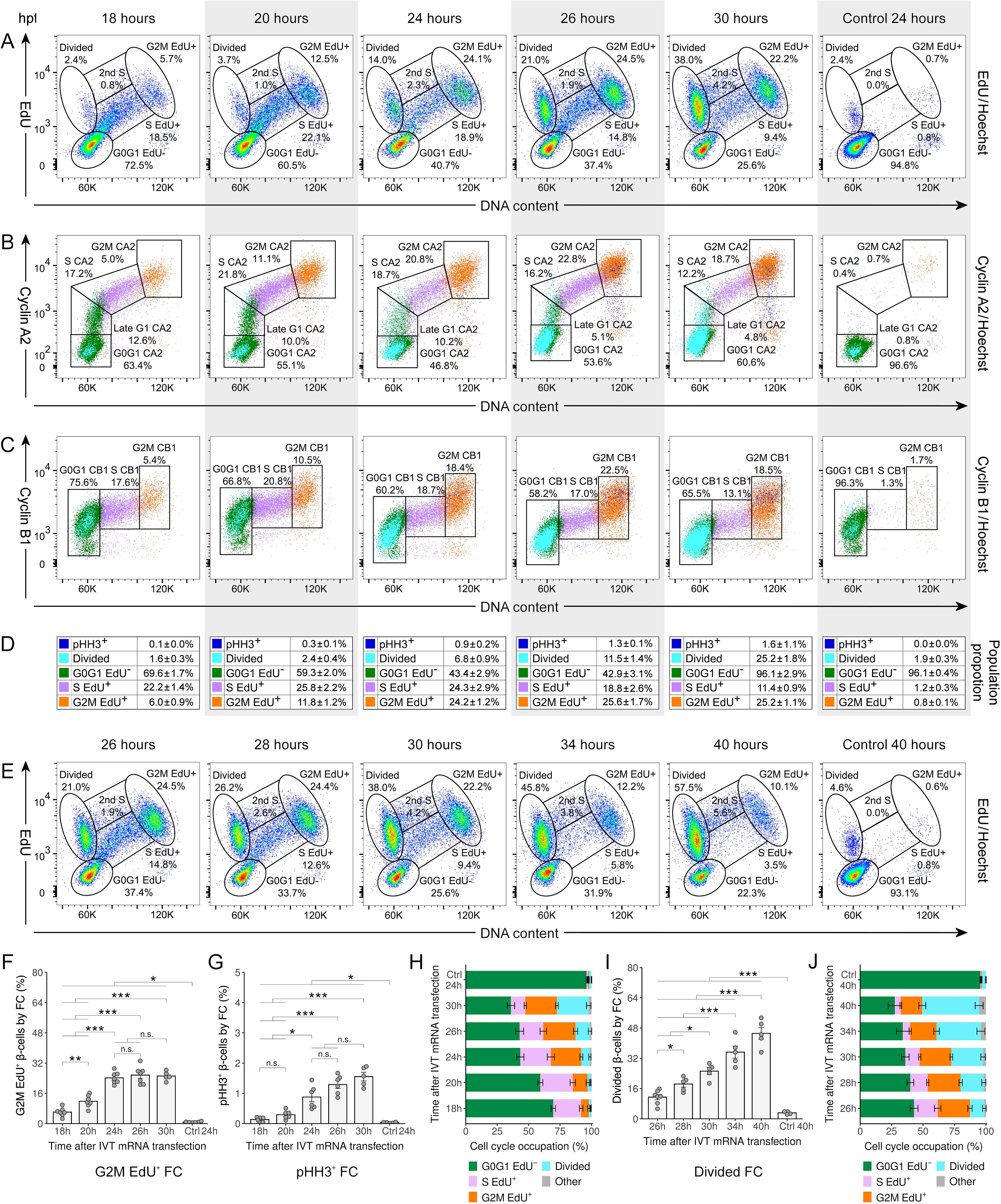
G2/M duration in β-cells is relatively uniform (≈ 7 hours) Representative flow cytometry plots illustrating G2/M progression and division into daughter cells in mRNA-treated and untreated β-cells at selected time points, highlighted by matching white and gray shading across panels (A–D). Cell populations are colour-coded in panels (B–D) and distribution graphs (H and J) as follows: G0/G1 EdU^-^(dark green), S EdU^+^ (violet), G2/M EdU^+^ (orange), divided β-cells (light blue), and pHH3^+^ (phosphorylated histone H3, dark blue). (A–C) Representative flow cytometry plots of β-cells showing EdU incorporation (A), Cyclin A2 (B), or Cyclin B1 expression (C) in combination with DNA content (see also Fig EV1). (D) Summary table showing percentages of β-cell populations derived from normalized EdU/DNA and pHH3/DNA flow cytometry analyses (mean ± SEM). (E) Representative flow cytometry plots demonstrating gradual appearance of divided β-cells visualized by EdU incorporation in combination with DNA content. (F) Proportion of G2/M EdU^+^ β-cells quantified from EdU/DNA flow cytometry (FC) plots and normalized to the original β-cell population (n=5–7 biological replicates). (G) Proportion of mitotic pHH3^+^ β-cells quantified from pHH3^+^/DNA flow cytometry plots (n=5–6 biological replicates). (H) Distribution of β-cells across cell-cycle phases derived from normalized EdU/DNA flow cytometry quantification shown in (A), with remaining cells grouped as “Other”. (I) Proportion of divided β-cells quantified from EdU/DNA plots and normalized to the original β-cell population (n=4–7 biological replicates). (J) Distribution of β-cells across cell-cycle phases derived from normalized EdU/DNA flow cytometry quantification shown in (E), with remaining cells grouped as “Other”. Bars represent mean ± SEM (F–J). Statistical significance was assessed using a linear mixed-effects model, including experiment as a random intercept and time as a categorical predictor (F, G, I), and allowing for group-specific residual variances (G). \*\*\**P* < 0.001, \*\**P* < 0.01, \**P* < 0.05, and n.s., not significant (Bonferroni-adjusted pairwise differences; omnibus test *P* < 0.001 for all tested models) Hpt, hours post-transfection

To independently assess S-phase progression, we performed immunofluorescence analysis using EdU and Cyclin A2 co-staining (Fig 3E, 3G, and 3I). IF analysis largely mirrored flow cytometry results, with both markers first detectable at 14 hpt and progressively increasing thereafter. From approximately 20 hpt onward, Cyclin A2 localisation became apparent in the cytoplasm, marking progression toward G2, as previously reported.(Silva Cascales *et al*, 2021) Taken together, while IF provided spatial information on Cyclin A2 localisation, early S-phase/late G1 events were more clearly detected by flow cytometry.

### G2/M duration in β-cells is relatively uniform (≈ 7 hours)

To gain insight into β-cell progression through G2/M, we analysed G2 and mitosis together, as mitosis is a very short phase that is difficult to resolve reliably in unsynchronised populations.(Chao *et al*, 2019) G2/M progression was mainly monitored by flow cytometry using EdU incorporation and DNA content. In addition, we also analysed Cyclin A2, Cyclin B1, and phosphorylated histone H3 at serine 28 (pHH3), a marker of chromatin condensation during mitosis (Goto *et al*, 1999; Pines & Hunter, 1991). G2/M duration was defined as the interval between entry into G2/M, characterised by maximal EdU intensity and 4N DNA content, and subsequent appearance within the divided population, characterised by 2N DNA content and persistent EdU positivity.

As described above, the earliest β-cells (group 1) were first detected in G2/M at 20 hpt (Fig 4A, 4F, and 4H). The proportion of G2/M β-cells increased rapidly over time, followed by cytokinesis and the appearance of a divided β-cell population (Fig 4A, 4D–4E, and 4I–4J). Detection of G2/M progression was further supported by increased Cyclin A2 and Cyclin B1 expression (Fig 4B and 4C), consistent with their known accumulation following completion of S-phase (Akopyan *et al*, 2014). In parallel, transient pHH3 positivity emerged from 20 hpt and peaked around 30 hpt, marking mitotic entry (Fig 4D and 4G).

The earliest significant fraction of divided β-cells (11.47 ± 1.38%) was detected at 26 hpt (Fig 4D and 4I). By 27 hpt, β-cell group 1 had completed division, corresponding to a G2/M duration of approximately 7 hours and a total cell-cycle length of approximately 26 hours (Table S1). Similar G2/M durations were observed for subsequent groups. Group 2 initiated G2/M at around 23 hpt and divided before 30 hpt, while group 3 entered G2/M before 27 hpt and divided before 34 hpt, corresponding to approximately 7-hour G2/M durations for both groups (Fig 4D, 4F, and 4I; Table EV1). In contrast, group 4 completed S-phase by 29 hpt but divided around 40 hpt, suggesting an apparently prolonged G2/M duration of up to 11 hours. However, this estimate was likely distorted by overlap with β-cells undergoing a second round of the cell cycle and cells exiting the cell cycle at later time points. Reliable estimation of G2/M duration was therefore not possible for group 5.

Overall, G2/M appeared to be the least variable cell-cycle phase, with an estimated duration of approximately 7 hours across the first three β-cell groups. Based on the overall results, we estimated total cell-cycle length at approximately 26 hours for group 1, 28 hours for group 2, and 30 hours for group 3. In contrast, group 4 displayed a longer apparent cell-cycle length of up to approximately 34 hours.

In non-stimulated, physiologically proliferating β-cells, the number of divided cells doubled between 40 and 70 hpt (Fig EV1), suggesting that their overall cell-cycle duration may be comparable to IVT-mRNA-stimulated β-cells. However, more detailed temporal analysis would be required for direct comparison.

### Gradual β-cell mitotic entry after synchronisation at the G2/M border

Next, we focused on the M-phase, a short period of the cell cycle that is difficult to capture in unsynchronised populations (Chao *et al*, 2019). To analyse mitotic progression quantitatively, we synchronised proliferating β-cells at the G2/M border before mitotic entry. For this purpose, we used RO-3306, a small-molecule CDK1 inhibitor that reversibly arrests cells in G2 before mitotic entry (Vassilev *et al*, 2006). This experimental design allowed accumulation of a large β-cell fraction in G2 while minimizing appearance of β-cells from S-phase after inhibitor removal. Proliferating islet cells were treated with RO-3306 in the presence of EdU. Following inhibitor washout, we released cells into fresh medium to enable synchronised progression through mitosis and cytokinesis, which we monitored until appearance of divided β-cells (Fig 5A).

**Fig 5:**
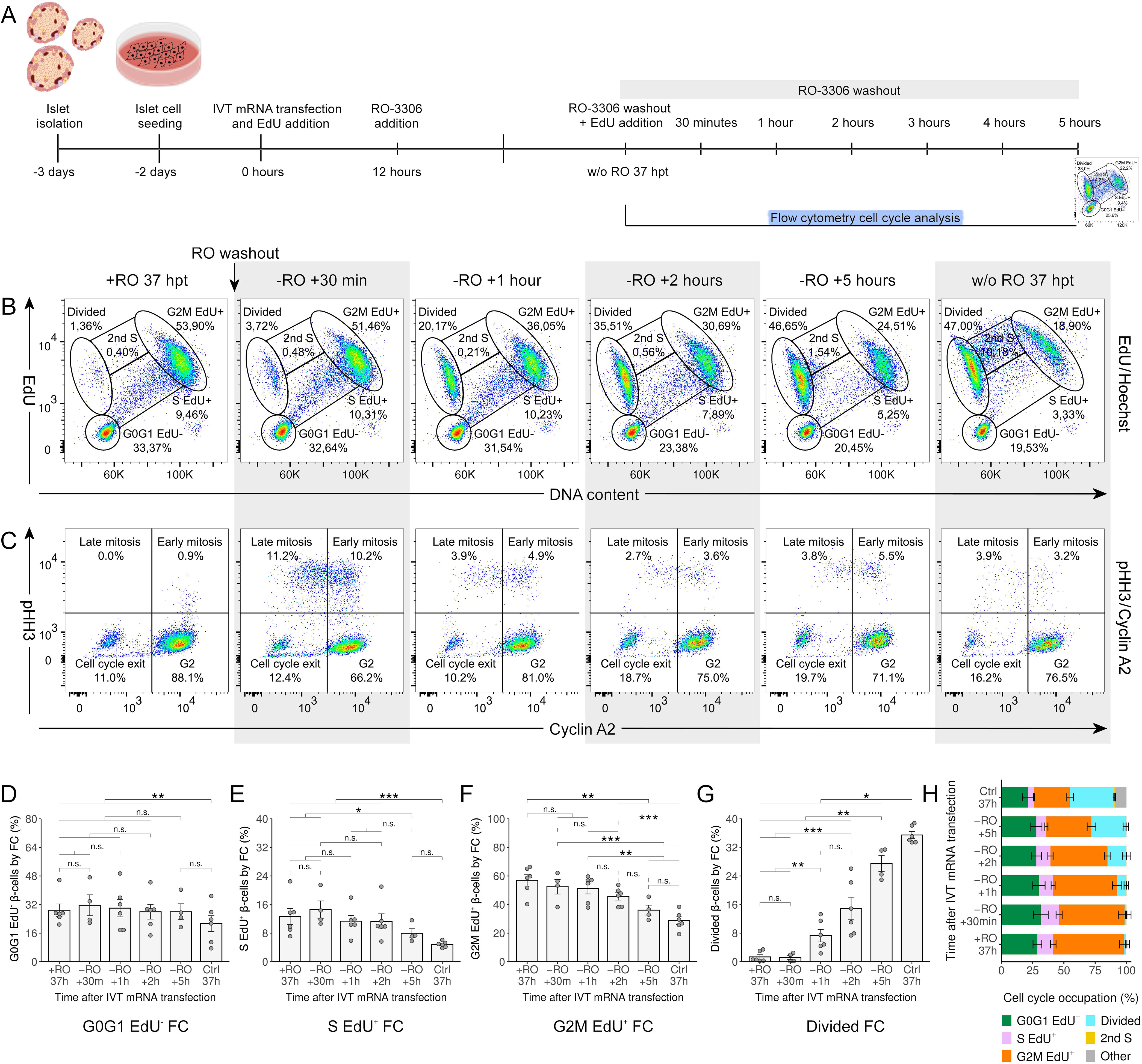
Gradual β-cell mitotic entry after synchronisation at the G2/M border (A) Schematic of mitosis tracking following G2/M arrest using the CDK1 inhibitor RO-3306 (see also Methods). (B–C) Representative flow cytometry plots of proliferating RO-treated and untreated β-cells showing EdU incorporation and DNA content (B) or phosphorylated histone H3 (pHH3) and Cyclin A2 expression within the EdU^+^ G2/M population (C) at selected time points (see also Fig EV4A and EV4B). The same time points are highlighted by white and gray shading across panels. (D–G) Proportions of G0/G1 EdU^-^(D), S EdU^+^ (E), G2/M EdU^+^ (F), and divided β-cells (G) quantified from EdU/DNA flow cytometry plots and normalized to the original β-cell population (n= 4–6 biological replicates) (H) Distribution of β-cells across cell-cycle phases derived from normalized EdU/DNA flow cytometry quantification, with remaining cells grouped as “Other”. Bars represent mean ± SEM (D–H). Statistical significance was assessed using a linear mixed-effects model, including experiment as a random intercept, time as a categorical predictor (D–G), and allowing for group-specific residual variances (G). \*\*\**P* < 0.001, \*\**P* < 0.01, \**P* < 0.05, and n.s., not significant (Bonferroni-adjusted pairwise differences; omnibus test *P* < 0.001 for all tested models). Hpt, hours post-transfection

RO-3306 treatment caused robust arrest of proliferating β-cells in G2 with high synchronisation efficiency (Fig 5B, 5F–5H, and EV4A). Thirty minutes after RO-3306 washout, we observed no significant increase in the fraction of divided β-cells, suggesting that mitosis required longer than 30 minutes. Concurrently, β-cells began to exhibit increased positivity for mitotic marker pHH3 (Fig 5C and EV4B). To further resolve mitotic stages, we co-analysed G2/M EdU^+^ β-cells for pHH3 and Cyclin A2, as Cyclin A2 degradation initiates during prometaphase (Den Elzen & Pines, 2001; Goto *et al*, 2002). One hour after RO-3306 removal, the first divided β-cells became detectable, consisting of 7.31 ± 1.75% of the original β-cell population (Fig 5G). After two hours, the fraction of divided β-cells approximately doubled to 14.91 ± 3.15%, indicating that mitosis and cytokinesis together require approximately 1-2 hours in the most rapidly dividing β-cells. The proportion of divided β-cells increased gradually over time and reached 27.48 ± 2.20% at 5 hours after RO-3306 washout compared to 35.48 ± 0.98% in the RO-3306-untreated sample.

The observed variability in mitotic activation likely reflects differences in the interval preceding mitotic entry rather than intrinsic heterogeneity in M-phase duration. This is supported by the gradual acquisition of pHH3 positivity after RO-3306 washout, indicating asynchronous entry into prophase (Fig 5C and EV4). Although most β-cells were expected to be arrested in G2 at the time of inhibitor removal, RO-3306 likely also partially inhibited CDK2, which regulates S-phase, and CDK4 which is needed for G1 (Vassilev *et al*, 2006). These effects could slow down whole cell-cycle progression and contribute to variability after RO-3306 washout. In addition, prolonged persistence in G2 may delay subsequent mitotic entry. This may result from extended CDK1 inhibition that hinders sustained accumulation and activation of mitotic regulators, including Cyclin B1, required for mitotic commitment (Akopyan *et al*, 2025). Consequently, after release from G2 arrest, β-cells may not enter mitosis in the same order as their earlier G1 entry timing.

Thus, partial synchronisation using RO-3306 enabled estimation of mitotic timing for the earliest-dividing β-cells, showing that M-phase lasted approximately 1–2 hours in this subpopulation. In contrast, mitotic entry of delayed β-cells could not be reliably determined under these conditions.

## DISCUSSION

The present study provides the first detailed characterization of cell-cycle progression in rat β-cells, revealing substantial heterogeneity in both cell-cycle entry and the duration of individual cell-cycle phases, particularly G1. IVT mRNA encoding cell-cycle regulators induced robust proliferation and enabled monitoring of complete cell-cycle progression in the majority of dividing β-cells over the three-day observation period. Establishing this high-resolution kinetic timeline allowed us to map distinct temporal dynamics before the onset of a second cell cycle that were previously unresolvable in primary islet cultures. Low proliferation rates reported in previous studies limited analysis of cell-cycle dynamics in primary β-cells and suggested uniform cell-cycle duration (Swenne, 1982; Hija *et al*, 2014; Saisho *et al*, 2009). In our study, the fastest-dividing β-cells completed the cell cycle in approximately 26 hours, whereas slower populations required up to 34 hours. Notably, our results revealed that longer cell-cycle duration was associated with later entry into G1. Earlier autoradiographic studies estimated the total cell-cycle length of fetal rat β-cells to 14.9 hours, partially due to a shorter G1 (2.5 hours) (Swenne, 1982). These differences likely stem from distinct analytical approaches and the fetal nature of the β-cells. Indeed, subsequent work has demonstrated that fetal β-cells differ substantially from mature β-cells in several functional aspects (Barsby & Otonkoski, 2022). Conversely, our estimates for S-phase and G2/M duration (both approximately 6 hours) are broadly consistent with this report. One explanation may be that the previous study predominantly analysed physiologically proliferating β-cells, which may exhibit a shorter G1.

Later studies examined the cell cycle of juvenile rat β-cells (approximately 40 hours) and adult mouse β-cells *in vivo* (approximately 27 hours post-stimulation) (Hija *et al*, 2014; Saisho *et al*, 2009). The latter study, which induced proliferation in only 4% of β-cells, is particularly relevant to our work, reporting durations of approximately 8, 5, 8, and 6 hours for the G0–G1 transition, G1, S, and G2/M, respectively. While our estimates for S-phase and G2/M durations are similar, our measurements indicate substantially longer G1 (13 hours for the fastest group 1), whereas the interval between cell-cycle stimulation and G1 entry was shorter (1 hour for the same group). The discrepancy may be explained by the choice of G1 marker, as Ki67, used in the previous study, is expressed following RB phosphorylation (Sobecki *et al*, 2017) and may delay detection of G1 events. In our parallel study, we also monitored Ki67 expression by IF, and detected its marked increase in later G1 (Koblas *et al*, 2026). In addition, glucokinase activation used in the previous study may promote β-cell proliferation through metabolic pathways, potentially altering the kinetics of cell-cycle entry compared to direct cell-cycle activation by CDK4–cyclin D.

Although IVT mRNA-mediated overexpression of CDK4–cyclin D1 represents a non-physiological stimulus, it provides an efficient and controllable tool to induce β-cell proliferation. Importantly, our results indicate that this stimulation did not substantially alter the intrinsic duration of individual cell-cycle phases compared to physiologically proliferating β-cells. In particular, the estimated lengths of S-phase and G2/M closely matched previous *in vivo* and *ex vivo* studies. Rather than dramatically accelerating the cell cycle, IVT mRNA-mediated stimulation primarily increases the fraction of β-cells that exit quiescence and enter the cell cycle. In this context, the gradual recruitment of β-cells into G1 likely reflects intrinsic heterogeneity in β-cell readiness to proliferate.

β-cells form a heterogeneous population with respect to their maturation status, including insulin expression, glucose responsiveness, and stress responses (Kobiita *et al*, 2023; Benninger & Hodson, 2018; Miranda *et al*, 2021). Less mature β-cells exhibit higher proliferative competence and respond more readily to metabolic (glucose, glucokinase) (Swenne, 1982; Hija *et al*, 2014), hormonal (prolactin or glucagon-like peptide) (Ackeifi *et al*, 2020, 1; Brelje *et al*, 1994) or small-molecule (Harmin) (Shen *et al*, 2015; Wang *et al*, 2019) stimulation. In contrast, mature β-cells typically require stronger activation of cell-cycle regulators to re-enter the cell cycle, such as ectopic overexpression of cell-cycle regulators in the form of IVT mRNA (Koblas *et al*, 2026) or viral carriers (Cozar-Castellano *et al*, 2004; Fiaschi-Taesch *et al*, 2009; Tiwari *et al*, 2015). Although CDK4, one of the main cell-cycle regulators, is physiologically expressed in β-cells, its endogenous level is low and insufficient to induce cell-cycle entry without additional stimulation (Fiaschi-Taesch *et al*, 2013a). Variability in expression of cell cycle-inhibitors may further contribute to differences in proliferative potential across β-cell subpopulations (Fiaschi-Taesch *et al*, 2013a, 2013b).

Considering the high efficiency of β-cell proliferation in our study, a large fraction of β-cells, including fully mature β-cells, were likely induced to enter the cell cycle. This underlies the heterogeneity in the timing of cell cycle-activation (G1 entry), ranging mostly from 1 hpt to 12 hpt, with the majority of β-cells entering G1 within the first 6 hpt. In addition to intrinsic β-cell properties, variability in transfection efficiency and resulting expression levels of cell cycle-regulators likely contributed to differences in G0–G1 transition kinetics. Consistent with this, control IVT GFP mRNA transfected into islet cells in our parallel study revealed substantial cell-to-cell variability in protein expression levels despite high overall β-cell transfection efficiency (approx. 80%) (Koblas *et al*, 2026). Importantly, heterogeneous timing of G1 entry has also been reported in established cell lines (Wang *et al*, 2017; Zetterberg & Larsson, 1985; Spencer *et al*, 2013), suggesting that variability in cell cycle-activation is not restricted to primary β-cells.

Both intrinsic β-cell heterogeneity and variability in transfection efficiency likely contributed to the pronounced variability in G1 length, which ranged from 13 to 19 hours and represented the most variable cell-cycle phase. During G1, cells grow and prepare for DNA replication, including progressive DNA licensing throughout the phase (Sosenko Piscitello *et al*, 2025; Fragkos *et al*, 2015; Bertoli *et al*, 2013). Consistent with previous observations, G1 duration following exit from quiescence can differ up to threefold among individual cells in cycling cell lines (Chao *et al*, 2019). In our study, however, G1 variability across β-cells was more limited (approximately 1.5-fold), which may reflect the generally prolonged G1 associated with exit from quiescence. One explanation for this extended G1 is the need to re-express components of the licensing machinery (Arias & Walter, 2007), whose expression is markedly reduced during quiescence (Kingsbury *et al*, 2005). Indeed, the first G1 following quiescence was reported to be significantly longer than subsequent G1 phases, reflecting a reduced number of licensed replication origins upon G0 exit (Matson *et al*, 2019). This mechanism may be relevant for naturally quiescent β-cells with distinct proliferative potential. The prolonged G1 observed in β-cells may therefore reflect the time required to restore sufficient replication licensing competence before S-phase entry. CDK4–cyclin D1 activity combined with relatively low but sufficient CDK2–cyclin E levels allowed prolonged G1 without premature acceleration of DNA replication (Koblas *et al*, 2026; Nussinov *et al*, 2024). From a cellular perspective, extending G1 may represent a protective strategy to ensure sufficient licensing and replication competence, thereby minimizing replication stress associated with an insufficient pool of dormant origins.

Within our study, we also compared flow cytometry and immunofluorescence as complementary approaches for cell-cycle analysis. Flow cytometry proved particularly useful for resolving subtle differences in the expression of weak cell-cycle markers, which could not be reliably distinguished by IF due to fluorescent background signal and subjective thresholding. Additionally, the combining EdU incorporation with DNA content measurement enabled accurate assignment of all cell-cycle phases, including divided cells, while allowing rapid analysis of large numbers of cells. In contrast, conventional IF does not directly provide information about DNA content, although quantitative image-based cytometry can partially address this limitation (Frölich *et al*, 2020a; Zerjatke *et al*, 2017). Nevertheless, IF offers a complementary advantage by enabling analysis of subcellular localisation of cell-cycle markers. For example, IF enables visualization of nuclear translocation of Cyclin B1 before mitotic entry (Pines & Hunter, 1991) or partial cytoplasmic translocation of Cyclin A2 in G2 (Silva Cascales *et al*, 2021). In addition, IF captures the cellular state immediately upon fixation, whereas flow cytometry involves additional processing before fixation that may affect transient cellular states.

With sufficiently dense temporal sampling, our approach is conceptually comparable to FUCCI-based systems that enable live-cell visualization of cell-cycle phases (Abe *et al*, 2013; Bajar *et al*, 2016). Nevertheless, unlike FUCCI, our strategy enables high-resolution reconstruction of cell-cycle progression in primary cells without the need for stable genetic reporters. The main limitation of our approach is the labour-intensive sampling at short intervals over several days.

While previous studies investigated the cell cycle of only a small β-cell subpopulations, here we provide a comprehensive characterisation of rat β-cell-cycle kinetics following proliferative stimulation. Our results demonstrate that proliferating β-cells form a heterogeneous population in which differences in G1 duration largely determine variability in overall cell-cycle length. In contrast, subsequent phases of the cell cycle exhibit relatively uniform timing. These findings highlight the central role of G1 regulation in controlling β-cell proliferation and suggest that the transition from quiescence represents a key bottleneck in β-cell-cycle entry. A better understanding of these regulatory mechanisms may help guide future strategies aimed at stimulating β-cell regeneration for diabetes therapy.

## METHODS

### EXPERIMENTAL MODEL AND STUDY PARTICIPANT DETAILS

#### Animal model

All experiments were performed with Islets of Langerhans cells isolated from male outbred Wistar rats aged 4-8 months (Charles River; Velaz). All animal procedures were approved by the Experimental Animal Welfare Committee of the Institute for Clinical and Experimental Medicine and the Ministry of Health of the Czech Republic (Permit Number: 30/2022; 53/2019). Animals were held according to the European Convention on Animal Protection and Guidelines on Research Animal Use in conventional breeding facility with 12/12 h light/dark cycle and free access to food and water.

#### Primary rat islet cells

Islets of Langerhans were isolated from pancreases of Wistar rats and dissociated into single-cell suspension. Islet cells were maintained on extracellular matrix coated plates under standard culture conditions.

#### HTB-9 cell line

Decellularized extracellular matrix produced by HTB-9 cells (American Type Culture Collection) was used to coat plates for islet cell culture. HTB-9 cells were maintained in RPMI 1640 (Thermo Fisher Scientific) supplemented with 10% heat-inactivated fetal bovine serum (FBS; Sigma-Aldrich).

### METHOD DETAILS

#### Generation of DNA templates for IVT transcription

Plasmids containing DNA template sequences were produced by GeneArt gene synthesis service (Thermo Fisher Scientific, Germany). Following plasmid amplification in E. coli HST08 strain (Takara Bio), plasmids were purified with CompactPrep Plasmid Midi Kit (Qiagen). Linearized DNA templates were amplified using PCR by means of Q5 Hot Start High-Fidelity DNA polymerase (New England Biolabs) and PCR products were purified with NucleoSpin Gel and PCR Clean-up Kit (Machery-Nagel). Sequence lengths were verified by 1.0% TBE agarose gel electrophoresis and DNA template concentration were assessed with Qubit dsDNA BR Assay Kit (Invitrogen). Both sequences were confirmed by Sanger sequencing and are provided in Tables S3 and S4.

#### IVT transcription

*In vitro* transcription (IVT) was performed as described previously (Koblas *et al*, 2026). PCR-amplified linear DNA templates were transcribed using MEGAscript T7 Transcription Kit (Invitrogen) for 3 hours at 37°C according to the manufacturer’s instructions. Additionally, 1-methylpseudouridine-5’-triphosphate (TriLink Biotechnologies) and 5’-methylcytidine-5′-triphosphate (TriLink Biotechnologies) were used to generate modified nucleoside-containing IVT mRNAs. For co-transcriptional capping of IVT mRNA 5’ ends was used CleanCap Reagent AG (TriLink Biotechnologies). IVT mRNA was then treated with 2 μl of Turbo DNase (Invitrogen) for 15 minutes at 37°C according to the manufacturer’s instructions. Antarctic Phosphatase (New England Biolabs) was used to remove non-capped 5′-triphosphate according to the manufacturer’s instructions. Additionally, USB Poly(A) Polymerase (Thermo Fisher Scientific) was used for polyadenylation of IVT mRNA 3’ ends. Following posttranscriptional modification, both IVT mRNAs were purified with MegaClear columns (Invitrogen) and dissolved in RNAsecure RNase Inactivation Reagent (Invitrogen). Final concentration of IVT mRNA was measured with Qubit RNA BR Assay Kit (Invitrogen), purity and integrity were assessed using Fragment Analyzer RNA Kit (Agilent) with a 5200 Fragment Analyzer System (Agilent). The prepared IVT mRNA encoding Cyclin D1 and CDK4 were stored at −80°C until use.

#### Pancreatic islet isolation

Islets were isolated according to a standardised procedure established in our laboratory (Kriz *et al*, 2012). Briefly, the pancreas was distended by intraductal injection of 15 mL collagenase (Collagenase Type V, Sigma-Aldrich) solution in Hankś Balanced Salt Solution (HBSS; 1.0 mg/mL, both from Sigma-Aldrich). The tissue was digested at 37 °C for 18 minutes until the pancreas dissociated into small fragments. The tissue was then filtered through a 500 µm mesh and purified using a discontinuous Ficoll density gradient (1.107 g/mL, 1.096 g/mL, 1.069 g/mL, 1.037 g/mL; Sigma-Aldrich). Isolated islets were handpicked under a stereomicroscope and cultured overnight in Connaught Medical Research Laboratories 1066 medium (CMRL-1066; PAN-Biotech) supplemented with 10% FBS (Sigma-Aldrich), 1x penicillin-streptomycin (100 U/mL penicillin and 100 μg/mL streptomycin; both from Sigma-Aldrich), 50 mM HEPES, and 1x GlutaMAX (Gibco), at 37 °C in a humidified atmosphere with 5% CO_2_. Residual exocrine tissue was removed by a second round of manual handpicking under the stereomicroscope the following day.

#### Islet cell culture and IVT mRNA transfection

Purified islets were dissociated using Accutase (approximately 1 μl/islet; Sigma-Aldrich) for 20 minutes at room temperature (RT) and plated onto wells coated with decellularized extracellular matrix produced by HTB-9 cells at a density of 70,000 cells per well. Islet cells were cultured in CMRL-1066 medium supplemented with 15% FBS (Sigma-Aldrich), 2x GlutaMAX, 2x MEM Non-Essential Amino Acids Solutions, 2 mM Sodium Pyruvate, 1x Insulin-Transferrin-Selenium, and 10 mM HEPES (all from Gibco). Transfection of islet cells with IVT mRNAs encoding Cyclin D1 and CDK4 was performed on day 3 after the seeding. Transfection pre-mix was prepared by mixing Lipofectamine MessengerMAX and Opti-MEM (both Invitrogen) at a ratio of 1:24 and incubated for 10 minutes at RT. Subsequently, IVT mRNAs were diluted in Opti-MEM to a concentration of 40 ng/μL. 80 ng of each mRNA was mixed with 10 µL per reaction of the pre-mix and incubated for 10 minutes at RT. Afterwards, the mixture was diluted with 40 µL of culture medium, and 50 µL was added per well. Cells were then pulsed with 5-Ethynyl-2′-deoxyuridine analog (EdU, Invitrogen) to a final concentration of 20 µM and maintained under standard culture conditions. The culture medium was refreshed after 48 hpt with fresh medium containing EdU.

#### Sample collection

For flow cytometry analysis, samples were collected at the following time points: 1 hpt, 2 hpt, and then every 2 hours for up to 34 hpt (the time point at 8 hpt was omitted due to redundancy in pilot experiments), and subsequently at 40 hpt, 50 hpt, 60 hpt, and 70 hpt (as shown in Fig 1A). Control samples included non-transfected islet cells at following time points: 0 hpt, 12 hpt, 24 hpt, 40 hpt, and 70 hpt. For sample collection, the culture medium including three PBS washes was collected. Cells were detached by incubation with pre-warmed TrypLE (37°C, Invitrogen) for 7 min and washed with HBSS. Then, cells were dissociated with Accutase for 20 minutes at RT, and fixed with 4% paraformaldehyde (PFA, Sigma-Aldrich) for 20 minutes on ice. Afterwards, cells were washed three times with 1% BSA in PBS, filtered through CellTricks disposable filters (50 μm pores, Sysmex) to obtain single-cell suspension. Finally, cells were fixed by drop-wise addition of ice-cold 70% ethanol (Hospital pharmacy) while vortexing and stored at −20°C until flow cytometry staining.

For immunofluorescence microscopy, samples between time points 0-24 hpt were fixed in 4% PFA for 20 minutes at RT, washed with PBS, incubated overnight in 1% PFA at 4°C, and processed for staining.

#### Cell-cycle synchronisation experiments

To synchronise islet cells in G2 phase, RO-3306 (20 µM, Tocris) was used. Starting at 12 hpt, islet cells were treated with RO-3306 for 24 hours, with an additional 20 µM dose administered 12 h after the initial treatment to maintain continuous CDK1 inhibition. At 37 hpt, the inhibitor was washed out four times with CMRL-1066 medium, after which islet cells were cultured in medium supplemented with 20 µM EdU for up to 5 hours. Samples were collected immediately before inhibitor removal, 30 minutes after washout, and subsequently at hourly intervals (Fig 5A). Control samples included proliferating cells cultured without the inhibitor and cells maintained in the continuous presence of the inhibitor without washout. All samples were processed for flow cytometry analysis as described above.

#### Flow cytometry and EdU labelling

Ethanol-fixed cells were washed three times with 1% BSA in PBS and permeabilized with 10x Permeabilization Buffer (1:10, Invitrogen) for 20 minutes at RT. Then, EdU was detected using Click-iT Plus EdU kit according to the manufacturerś instructions (Invitrogen). Briefly, cells were incubated with Click-iT Plus reaction cocktail (containing Click-it Reaction Buffer, Copper Protectant, Alexa Fluor 488 picolyl azide, and Reaction Buffer Additive) for 30 minutes at RT. From this step onward, all incubations were performed in the dark. Cells were washed three times with 1x Permeabilization Buffer and blocked with 2% rat (Abcam) and donkey sera (Jackson ImmunoResearch) in 1x Permeabilization Buffer for 15 minutes at RT. Samples were then incubated for 1 hour at RT with a mouse anti-Insulin antibody (1:200, Sigma-Aldrich) and one of the following rabbit antibodies: anti-phospho-Retinoblastoma protein (Ser807/811, 1:400, Cell Signaling Technology), anti-Stem Loop Binding Protein (1:100, Novus), anti-Cyclin A2 (1:400, Abcam) or anti-Cyclin B1 (1:300, Abcam), all diluted in block solution. After three washes with 1x Permeabilization Buffer, cells were incubated with secondary antibodies, anti-mouse BV 711 (1:100, BD Biosciences) and anti-rabbit Alexa Fluor 647 plus (1:100, Invitrogen) diluted in 1x Permeabilization Buffer for 30 minutes at RT. Cells were then washed three times with 1x Permeabilization Buffer, blocked again with 2% Rabbit serum in 1x Permeabilization Buffer for 15 minutes, and incubated with rabbit anti-phospho-Histone H3 (Ser28) conjugated to Alexa Fluor 555 (1:450, Abcam) for 30 minutes at RT. Finally, cells were washed three times and stained with Hoechst 33342 (1:4000, Invitrogen) until flow cytometry analysis. Samples were analysed using FACSymphony flow cytometer (BD Biosciences). In addition to the forward scatter and side scatter, the following fluorescence channels were detected: Hoechst 33342 (355 nm laser: 450/50), Alexa Fluor488 (488 nm laser: 530/30), Alexa Fluor 555 (561 nm laser: 586/15), Alexa Fluor 647 (637 nm laser: 670/30), and BV711 (405 nm laser: 710/50). Data were processed in BD FlowJo v10.8.1 software (BD Biosciences). Cell doublets and debris were excluded from all analyses based on forward and side scatter properties.

#### Immunofluorescence microscopy

Fixed islet cells with PFA were washed three times with PBS and permeabilized with 0.1% Triton X-100 (Sigma-Aldrich) in PBS for 20 minutes at RT. After three additional PBS washes, EdU was detected using the Click-iT Plus EdU kit as described above (Invitrogen). All subsequent steps were performed in the dark. Cells were blocked with 5% donkey serum in blocking solution containing 0.05% Triton X-100, 0.75% Glycine, and PBS for 30 minutes at RT. Cells were then incubated for 1 hour with mouse anti-Insulin primary antibody (1:200, Sigma-Aldrich), together with either rabbit anti-phospho-Retinoblastoma protein (Ser807/811, 1:400, Cell Signaling) or rabbit anti-Cyclin A2 (1:400, Abcam), diluted in antibody diluent solution (0.2% cold fish skin gelatin, 0.01% sodium azide, 0.5% Triton X-100, and 1% BSA and PBS). After three PBS washes, cells were incubated for 1 hour at RT with secondary donkey antibodies conjugated to anti-mouse BV 711 (1:500, BD Biosciences) or anti-rabbit Alexa Fluor 647 plus (1:500, Invitrogen) in blocking solution. Following three PBS washes, nuclei were counterstained with NucBlue Fixed Cell Stain ReadyProbes reagent (2 drops/mL, Invitrogen) diluted in PBS for 5 minutes at RT, followed by six PBS washes. Finally, cells were mounted in Dabco Mowiol mounting medium (2.5% 1,4-diazabicyclo-[2,2,2]-octane, 4.8% Mowiol, 12% glycerol, Carl Roth, and 0.2 M Tris HCl). Immunofluorescence images were acquired using an EVOS M7000 Imaging System (Invitrogen). For quantification of individual β-cells in individual cell-cycle phases, at least 20,000 cells per well were analysed in merged DAPI, Insulin, EdU, and either Cyclin A2 or pRB images using Celleste Image Analysis Software version 6 (Invitrogen).

### QUANTIFICATION AND STATISTICAL ANALYSIS

#### Recalculation of β-cell population frequencies from EdU/DNA plots

To estimate β-cell proportions in individual cell-cycle phases normalized to the original β-cell population at 0 hpt, values obtained from EdU/DNA flow cytometry plots were recalculated to account for cell division. The total number of original β-cells was defined as:

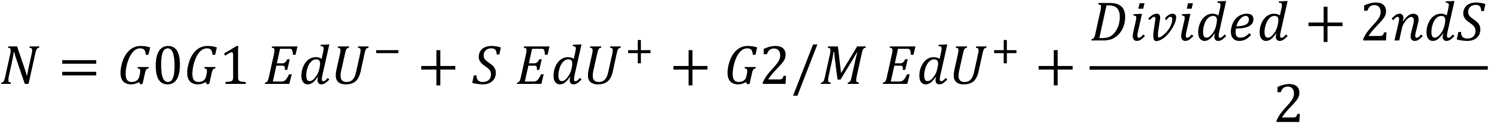

where G0/G1 EdU^-^, S EdU^+^, G2/M EdU^+^ represent percentage of β-cells in the respective cell-cycle phase identified by EdU/DNA flow cytometry analysis, while Divided and 2ndS correspond to β-cells that completed division and undergo a second G0 or G1 phase or entered a second round of S phase.

The proportion of β-cells in each population relative to the original β-cell number was calculated for G0/G1 EdU^-^β-cells as:

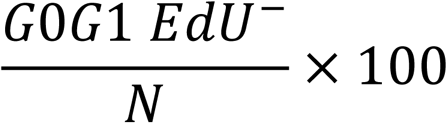

S EdU^+^ β-cells as:

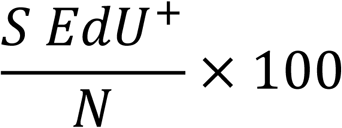

G2/M EdU^+^ β-cells as:

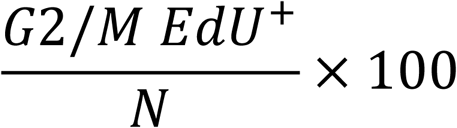

Divided β-cells as:

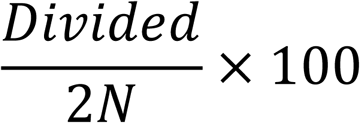

2nd S β-cells as:

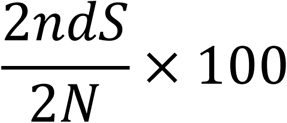

Determination of individual β-cell cell-cycle phase length

Lengths of individual cell-cycle phases were calculated as the difference between the time point at which a given group of β-cells entered a specific phase and time point at which the same group entered the subsequent phase. For values derived from normalized EdU/DNA flow cytometry plots (all phases except G1), the proportion of β-cells in a given phase was inferred from the corresponding fraction including subsequent phases, to account for potential differences in progression speed. All derived values were normalized to non-mRNA-treated control samples. If a given group of β-cells transitioned to the subsequent phase between the two 2-hour sampling intervals, the interval was subdivided into quarters to estimate phase lengths with 1-hour resolution. All flow cytometry-derived values normalized to the original number of β-cells in case of EdU are reported in Table S1 and calculated phase lengths are reported in Table S2.

#### Statistical analysis

All statistical analyses were performed using R, version 4.4.3.(R Development Core Team (2008). Data were analysed using linear mixed-effects models implemented in the glmmTMB package (Brooks *et al*, 2017).

The proportion of β-cells classified into the given cell population was used as the outcome variable, modeled with a Gaussian error distribution; separate models were fitted for each cell population. Experimental group (time point or untreated control, where applicable) was used as a categorical fixed-effect predictor. Each independent experiment (individual islet isolation) was included as a random intercept to reflect the dependency of observations within a single experiment. Overall effects of the experimental group (omnibus tests) were assessed using Wald chi-square tests. Model assumptions were examined using simulated randomised quantile residuals (via the DHARMa package). A model with constant residual variance was used as the default; however, group-specific residual variances (modeled via a dispersion formula) were introduced when their inclusion (i) was supported by a visual pattern of residual variance changing with the group mean, (ii) decreased the Akaike information criterion (AIC), and (iii) improved residual diagnostics. For post hoc analysis, groups were compared pairwise using the emmeans package (Lenth *et al*, 2026). All pairwise p-values and confidence intervals were adjusted using the Bonferroni correction, with statistical significance defined as a two-sided P < 0.05. Data are presented as mean ± SEM.

The number of independent experiments (n) performed for each condition is specified in the corresponding Figure legends; n represents the number of independent biological replicates, each corresponding to an individual islet isolation. Sample sizes were determined based on experimental feasibility; no formal power analysis was performed. Given the nature of the *in vitro* time-course experiments, randomisation and blinding were not applicable. No data, experiments, or observations were excluded from the analyses.

Details of model diagnostics, 95% confidence intervals for pairwise differences, and statistical code are available online at https://filip-tichanek.github.io/beta_cells/ and archived on Zenodo at https://doi.org/10.5281/zenodo.22109481.

### KEY RESOURCES TABLE

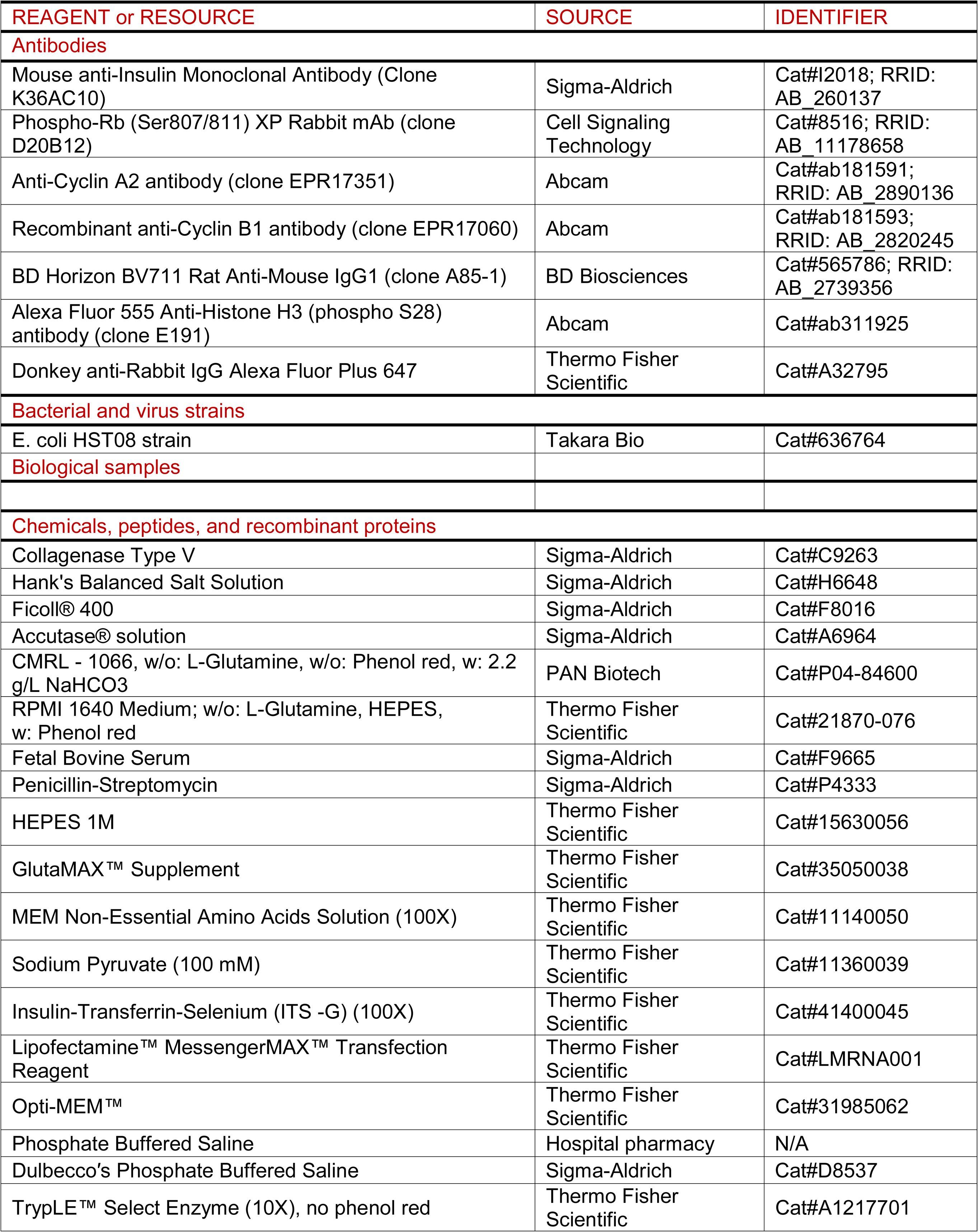
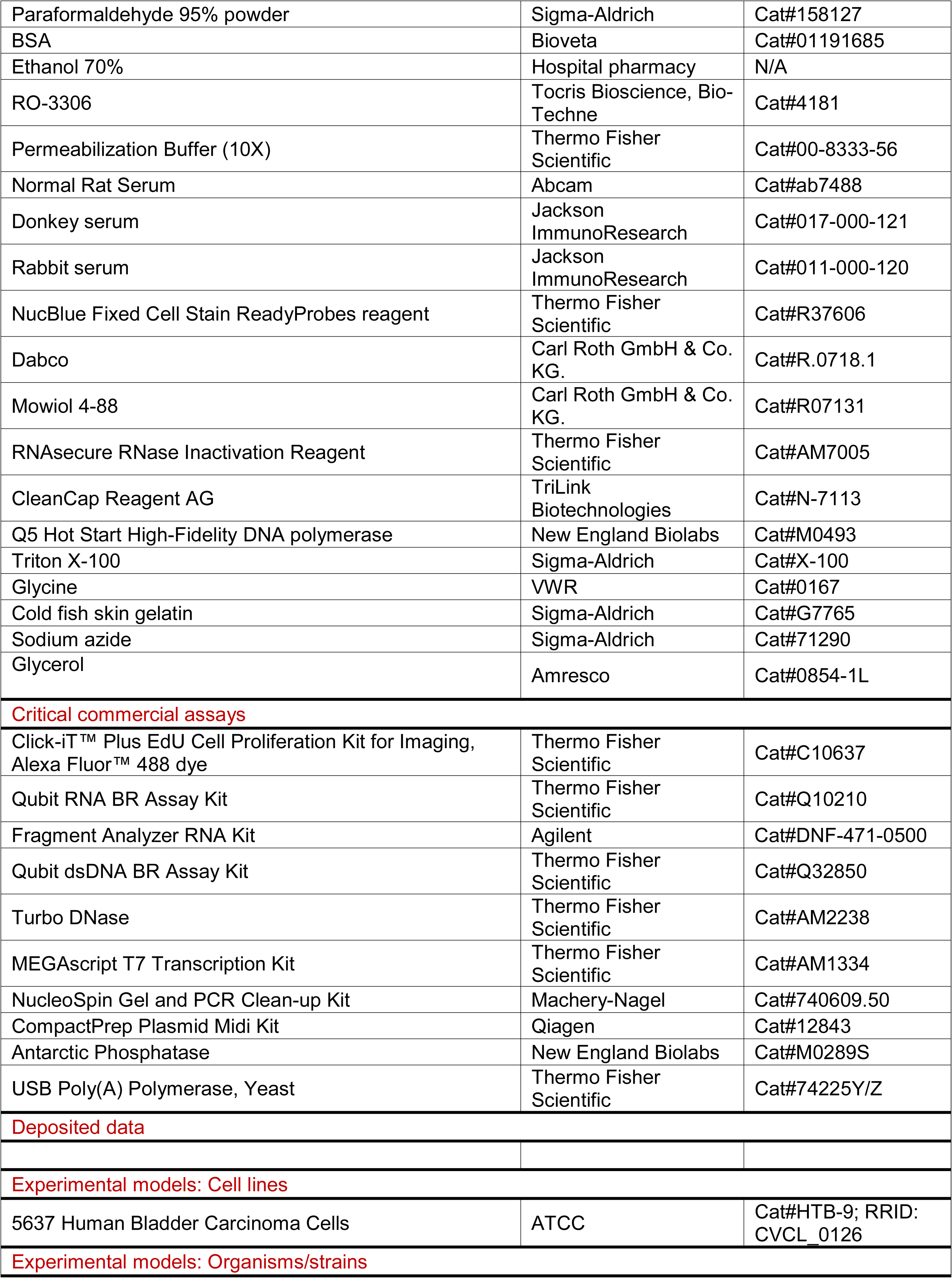
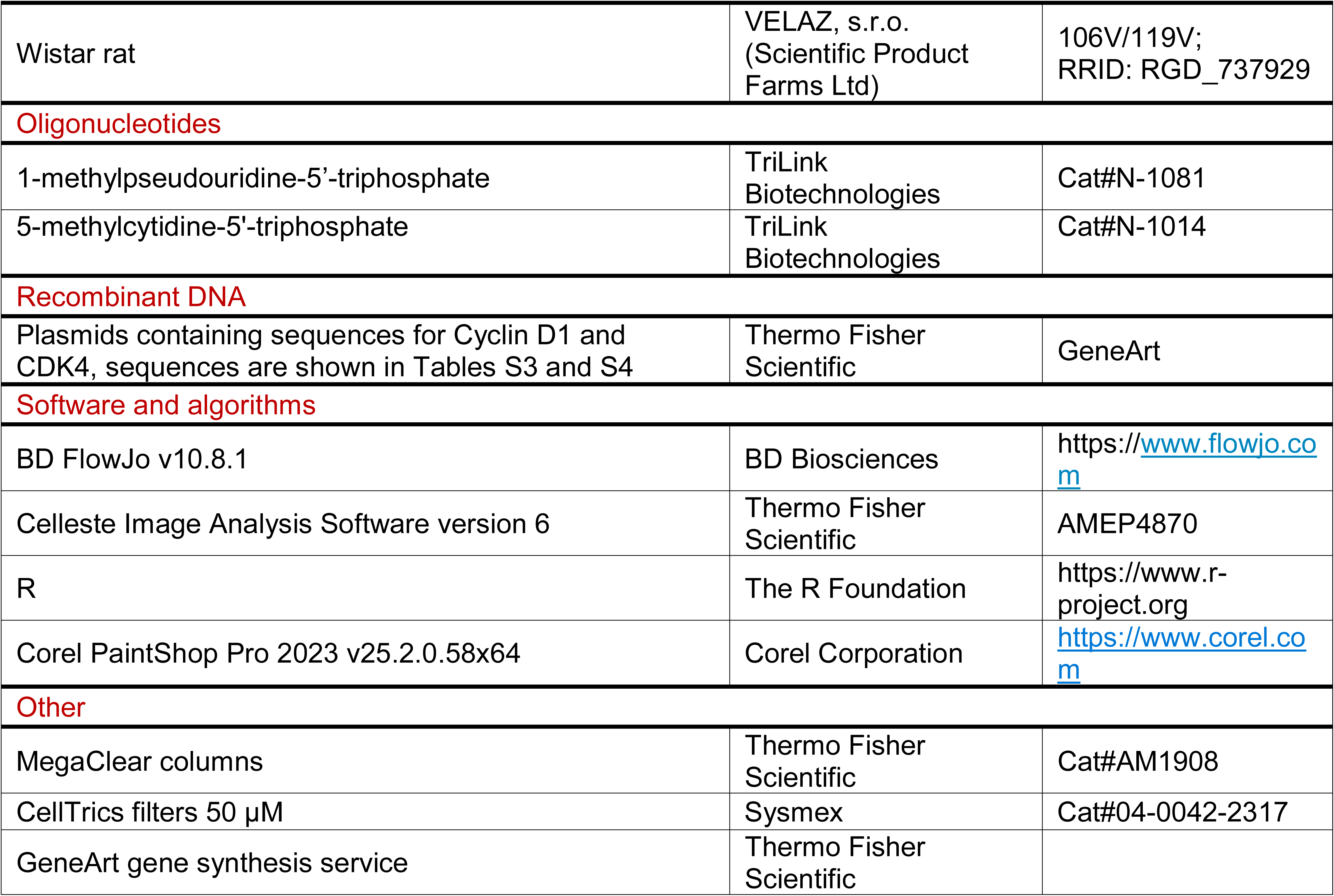

## RESOURCE AVAILABILITY

### Lead contact

Further information and requests for resources and reagents should be directed to and will be fulfilled by the lead contact, Tomas Koblas.

### Materials availability

Plasmids generated from this study are available from the lead contact upon request, other materials used in this study are commercially available.

### Data and code availability

Processed data from flow cytometry are provided in Table S1. The rendered statistical report, including model diagnostics and statistical test outputs, is available at https://filip-tichanek.github.io/beta_cells/. The underlying R code and processed analysis data are publicly available at https://github.com/filip-tichanek/beta_cells/ and archived on Zenodo at https://doi.org/10.5281/zenodo.22109481 (Bittenglova *et al*, 2026). Any additional information required to reanalyse the data reported in this paper is available from the lead contact upon request.

## ACKNOWLEDGMENTS

Supported by the Ministry of Health of the Czech Republic in cooperation with the Czech Health Research Council within the National Institute CarDia, project No. NW26A-CARDIA (J.K. and F.S.). Additional support was provided by: Ministry of Health, Czech Republic – conceptual development of research organisation („Institute for Clinical and Experimental Medicine–IKEM, IN 00023001“) (F.S.), by Charles University, project GA UK No. 299122 (K.B.), by Czech Science Foundation project number 24-11364S/26-11364S (T.K.), and by Czech Science Foundation, project number 25-15876S (F.S.). We thank Magdalena Spitalnikova Vertatova for assistance with microscopy and sample collection.

## AUTHOR CONTRIBUTIONS

K.B. and T.K. conceived the study and designed experiments. K.B., K.Z., I.L. and T.K. performed experiments. K.Z. conducted animal experiments. K.B. and F.T. analysed data. F.T. performed statistical analysis, K.Z., F.S., J.K., and T.K. reviewed and edited the manuscript. F.S., J.K., and T.K. supervised the study. K.B. prepared Figure and wrote the manuscript.

## DECLARATION OF GENERATIVE AI AND AI-ASSISTED TECHNOLOGIES IN THE WRITING PROCESS

During the preparation of this work, the authors used Chat GPT and Gemini in order to language correction and improvement of readability. After using this tool or service, the authors reviewed and edited the content as needed and take full responsibility for the content of the publication.

## EXPANDED VIEW FIGURE LEGENDS

**Table EV1:** Values derived from pRB/DNA and EdU/DNA flow cytometry plots and normalised to the original number of β-cells at all time points. Values are represented as means (n=4–7 biological replicates depending on time points)

|  | G1 pRB | S EdU<sup>+</sup> | G2/M EdU<sup>+</sup> | Divided | 2nd S |
| --- | --- | --- | --- | --- | --- |
| 0 hpt Control | 2.47 |  |  |  |  |
| 1 hpt | 15.79 |  |  |  |  |
| 2 hpt | 26.17 |  |  |  |  |
| 4 hpt | 40.89 |  |  |  |  |
| 6 hpt | 50.18 |  |  |  |  |
| 10 hpt | 60.24 |  |  |  |  |
| 12 hpt | 64.54 | 3.01 | 0.76 | 0.47 | 0.04 |
| 12 hpt Control | 2.82 | 0.81 | 0.66 | 0.29 | 0.01 |
| 14 hpt | 55.00 | 10.99 | 1.48 | 1.04 | 0.19 |
| 16 hpt | 49.95 | 16.39 | 2.69 | 1.20 | 0.28 |
| 18 hpt | 46.62 | 22.17 | 6.07 | 1.62 | 0.47 |
| 20 hpt | 38.57 | 25.85 | 11.78 | 2.40 | 0.65 |
| 22 hpt | 28.95 | 24.85 | 19.34 | 3.97 | 1.03 |
| 24 hpt | 28.75 | 24.37 | 24.16 | 6.75 | 1.28 |
| 24 hpt Control | 2.47 | 1.17 | 0.81 | 1.88 | 0.02 |
| 26 hpt |  | 18.98 | 25.61 | 11.34 | 1.19 |
| 28 hpt |  | 13.84 | 26.22 | 18.27 | 1.66 |
| 30 hpt |  | 11.41 | 25.17 | 25.08 | 2.19 |
| 32 hpt |  | 10.51 | 24.01 | 29.01 | 2.44 |
| 34 hpt |  | 5.95 | 20.81 | 34.91 | 3.86 |
| 40 hpt |  | 4.70 | 17.10 | 44.86 | 5.28 |
| 40 hpt Control |  | 0.60 | 0.60 | 3.17 | 0.03 |
| 50 hpt |  | 1.93 | 18.37 | 55.56 | 3.86 |
| 60 hpt |  | 0.94 | 9.55 | 63.34 | 0.94 |
| 70 hpt |  | 1.00 | 12.37 | 68.22 | 0.96 |
| 70 hpt Control |  | 1.05 | 0.71 | 6.15 | 0.02 |

**Table EV2:**
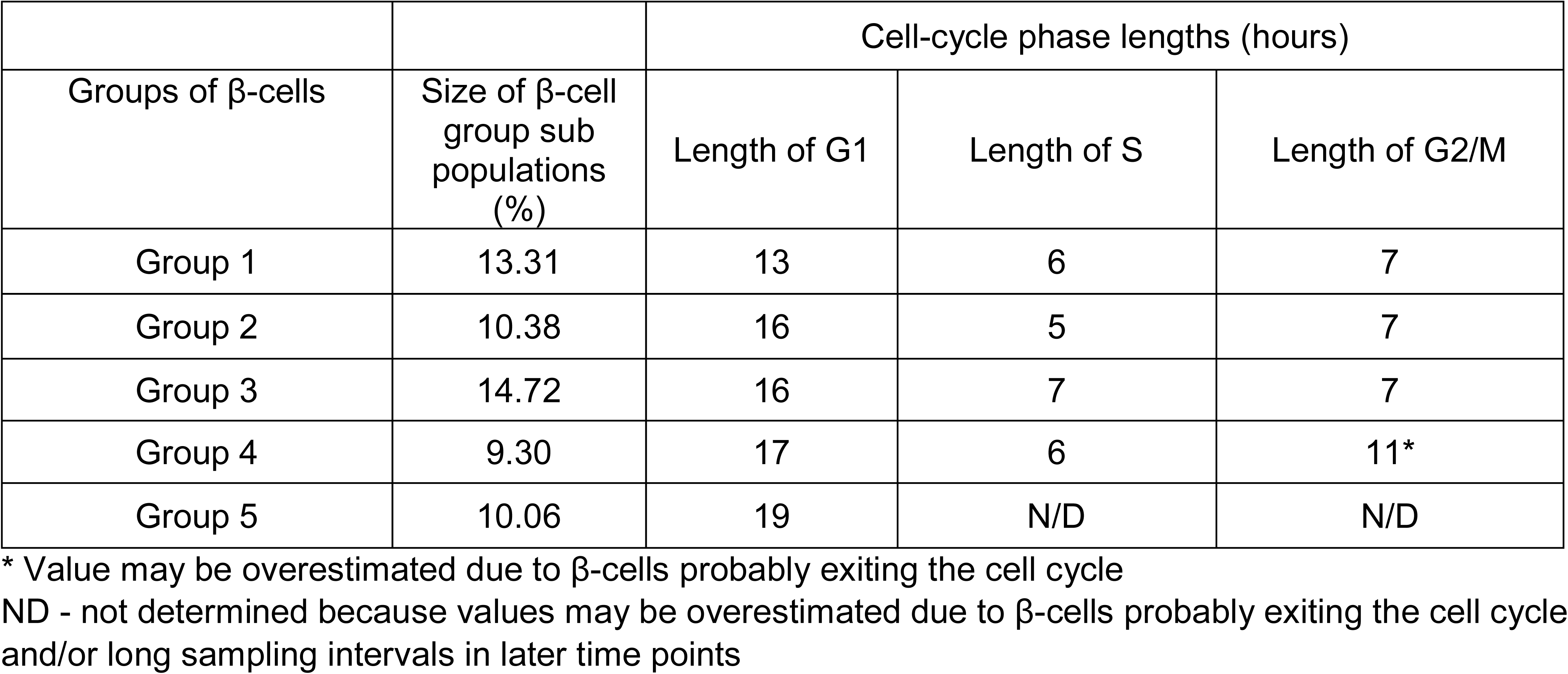
Table summarizing sizes of β-cell group and lengths of individual cell-cycle phases.

| Groups of β-cells | Size of β-cell group sub populations (%) | Cell-cycle phase lengths (hours) | | |
| --- | --- | --- | --- | --- |
|  |  | Length of G1 | Length of S | Length of G2/M |
| Group 1 | 13.31 | 13 | 6 | 7 |
| Group 2 | 10.38 | 16 | 5 | 7 |
| Group 3 | 14.72 | 16 | 7 | 7 |
| Group 4 | 9.30 | 17 | 6 | 11* |
| Group 5 | 10.06 | 19 | N/D | N/D |

\* Value may be overestimated due to β-cells probably exiting the cell cycle
ND - not determined because values may be overestimated due to β-cells probably exiting the cell cycle and/or long sampling intervals in later time points

Figure EV1. Half of β-cells complete division within 40 hours after IVT mRNA-induced cell-cycle activation, related to Figure 1

A) Representative flow cytometry plots of mRNA-treated and untreated β-cells showing EdU incorporation and DNA content at all time points.

Hpt, hours post-transfection

Figure EV2: β-cell S-phase duration is independent of the timing of cell-cycle entry, related to Figure 3

A) Representative flow cytometry plots of mRNA-treated and untreated β-cells showing Cyclin A2 (CA2) expression and DNA content at later time points. Note the proportion of proliferating Cyclin A2^-^G2/M β-cells (in plots labelled “CC exit/mitosis CA2^-^) increased over time, suggesting possible progression towards the cell-cycle exit.

Hpt, hours post-transfection

Figure EV3: β-cell G1 duration varies with the timing of G1 entry, related to Figure 2

(A) Representative flow cytometry plots of mRNA-treated and untreated β-cells showing RB phosphorylation (pRB) and DNA content at all analysed time points. Note in the later time points, the percentage of proliferating pRB^-^G2/M β-cells (in plots labelled “CC exit/mitosis RB^-^) increased over time, suggesting possible progression towards the cell-cycle exit.

Hpt, hours post-transfection

Figure EV4: Gradual β-cell mitotic entry after synchronisation at the G2/M border, related to Figure 5

(A–B) Representative flow cytometry plots of proliferating RO-treated β-cells showing EdU incorporation in combination with DNA content (A) and histone H3 phosphorylation (pHH3) in combination with Cyclin A2 expression (B) at all timepoints after RO-3306 washout.

Figure EV5: Representative gating strategies from scatter plots to EdU versus DNA content and additional markers in insulin positive cells

(A–E): Representative gating strategies for RB phosphorylation (A), Cyclin A2 expression (B), SLBP expression (C), Cyclin B1 expression (D) versus DNA content, and for histone H3 phosphorylation versus Cyclin A2 expression in the G2/M EdU^+^ population (E).

**Table EV3:**
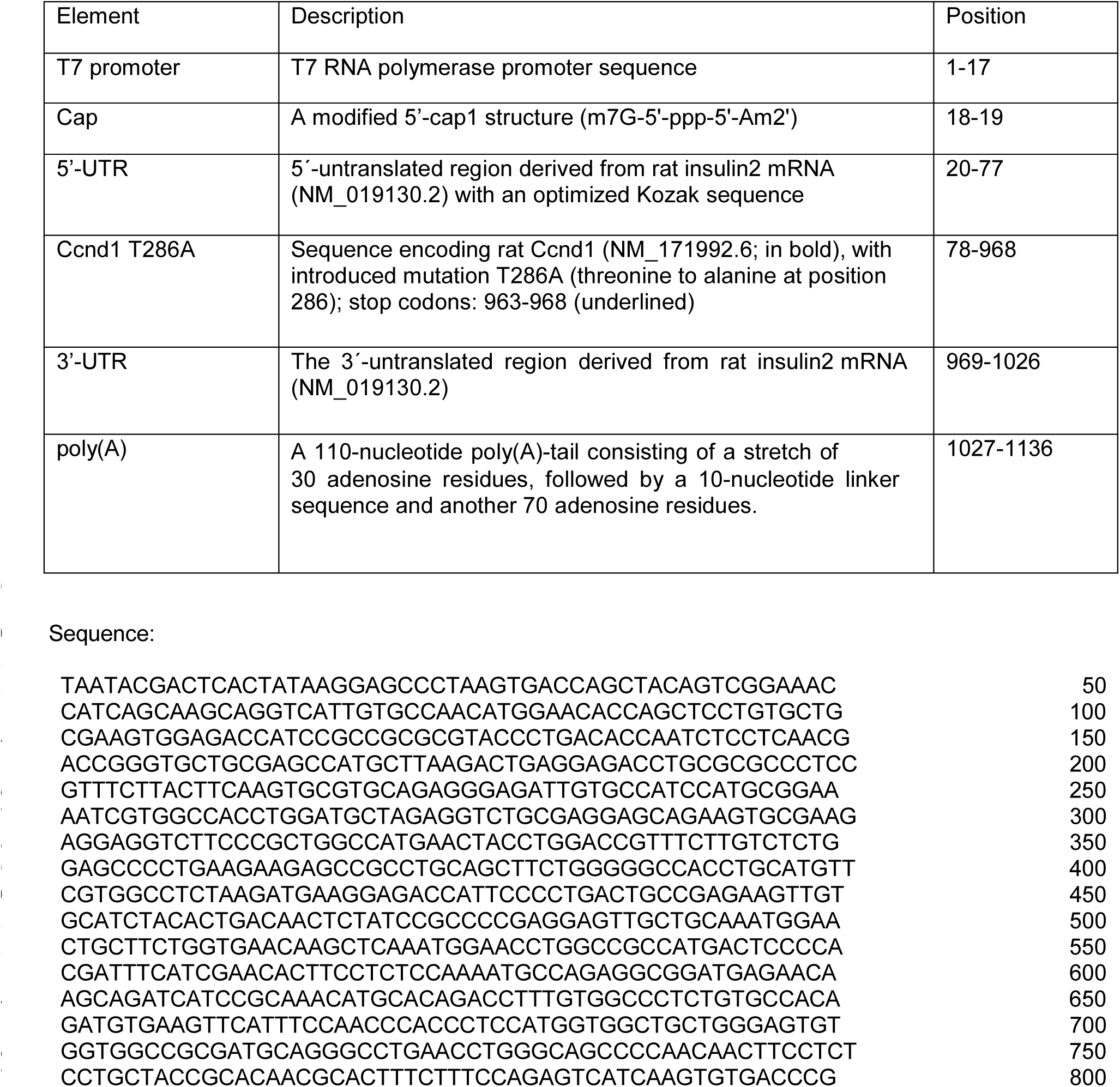
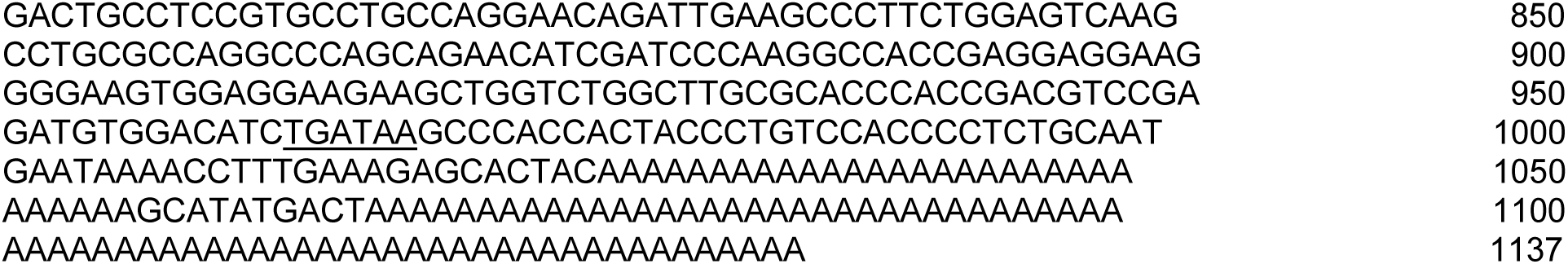
DNA template sequence encoding rat Ccnd1 T286A IVT mRNA.

**Table EV4:** DNA template sequence encoding rat CDK4 IVT mRNA.

| Element | Description | Position |
| --- | --- | --- |
| T7 promoter | T7 RNA polymerase promoter sequence | 1-17 |
| Cap | A modified 5'-cap1 structure (m7G-5'-ppp-5'-Am2') | 18-19 |
| 5'-UTR | 5'-untranslated region derived from rat insulin2 mRNA (NM_019130.2) with an optimized Kozak sequence | 20-77 |
| Cdk4 | Sequence encoding rat Cdk4 (NM_053593.3; in bold); stop codons: 987-992 (underlined) | 78-992 |
| 3'-UTR | The 3'-untranslated region derived from rat insulin2 mRNA (NM_019130.2) | 993-1050 |
| poly(A) | A 110-nucleotide poly(A)-tail consisting of a stretch of 30 adenosine residues, followed by a 10-nucleotide linker sequence and another 70 adenosine residues. | 1051-1160 |

Sequence:
AAAAAAAAAAAAAAAAAAAAAAAAAAAAAAAAAAAAAAAAAAAAAAAAAAAAAA 1150
AAAAAAAAAA 1160

